# A CTDNEP1-lipin 1-mTOR regulatory network restricts ER membrane biogenesis to enable chromosome motions necessary for mitotic fidelity

**DOI:** 10.1101/2021.03.02.433553

**Authors:** Holly Merta, Jake W. Carrasquillo Rodríguez, Maya I. Anjur-Dietrich, Mitchell E. Granade, Tevis Vitale, Thurl E. Harris, Daniel J. Needleman, Shirin Bahmanyar

## Abstract

The endoplasmic reticulum (ER) dramatically restructures in open mitosis to become excluded from the mitotic spindle; however, the significance of ER reorganization to mitotic progression is not known. Here, we demonstrate that limiting ER membrane biogenesis enables mitotic chromosome movements necessary for chromosome biorientation and prevention of micronuclei formation. Aberrantly expanded ER membranes increase the effective viscosity of the mitotic cytoplasm to physically restrict chromosome dynamics – slowed chromosome motions impede correction of mitotic errors induced by transient spindle disassembly, leading to severe micronucleation. We define the mechanistic link between regulation of ER membrane biogenesis and mitotic fidelity by demonstrating that a CTDNEP1-lipin 1-mTOR regulatory network limits ER lipid synthesis to prevent chromosome missegregation. Together, this work shows that ER membranes reorganize in mitosis to enable chromosome movements necessary for mitotic error correction and reveal dysregulated lipid metabolism as a potential source of aneuploidy in cancer cells.

## Introduction

The endoplasmic reticulum (ER) is a large membrane-bound organelle composed of interconnected membrane sheets and tubules that are continuous with the nuclear envelope (Baumann and Walz, 2001; Friedman and Voeltz, 2011). In animal cells, ER membranes are spread throughout the cytoplasm extending from the NE along the microtubule (MT) cytoskeleton to the cell periphery (English and Voeltz, 2013). The ER loses most of its interactions with MTs in mitosis (Smyth et al., 2012; Wang et al., 2013), and NE/ER membranes undergo extensive remodeling to become excluded from the region occupied by the mitotic spindle in metaphase (Champion et al., 2017; Liu and Pellman, 2020; Lu et al., 2011; Puhka et al., 2012; Puhka et al., 2007). ER membranes remain excluded from mitotic chromosomes and the region where spindle midzone microtubules assemble during chromosome segregation. After mitotic exit, and coincident with mitotic spindle disassembly, ER membranes and associated integral NE proteins gain access to decondensing chromosomes to reform the NE (Liu and Pellman, 2020). The mechanisms that drive ER membrane exclusion from spindle MTs throughout mitosis are not fully understood; however, both active mechanisms, mediated by the minus-end MT motor dynein (Turgay et al., 2014) and by the ER tubule shaping proteins REEP3/4 (Kumar et al., 2019; Schlaitz et al., 2013), and passive mechanisms, resulting from loss of MT-ER interactions, are thought to be involved (Liu and Pellman, 2020).

Little is known about the significance of the spatial reorganization of ER membranes in mitosis. The remodeling of ER membranes and their location away from the mitotic spindle may facilitate equal partitioning of the ER to daughter cells (Champion et al., 2017). Some evidence also suggests that the ER surrounding spindle MTs serves as part of an organelle-exclusion ‘spindle envelope’ that spatially confines mitotic proteins to the spindle region (Schweizer et al., 2015). Persistent ER membrane contacts with mitotic chromosomes, but not with spindle MTs, have been associated with higher incidences of chromosome missegregation (Champion et al., 2019; Luithle et al., 2020; Schlaitz et al., 2013; Smyth et al., 2012). These studies suggest that the physical removal of ER membranes from mitotic chromosomes is necessary for accurate chromosome segregation; however, how the spatial reorganization of ER membranes and its clearance from the spindle region contributes to mitotic fidelity remains unknown.

The need for ER membranes to be excluded from the spindle region may be met by restricting the production of ER membranes prior to entry into mitosis. Membranes double in S-phase (Jackowski, 1994), but whether ER lipid synthesis is coordinated with mitotic reorganization of the ER to ensure membrane clearance and mitotic fidelity has not been tested. A key enzyme required for ER membrane biogenesis is the Mg^2+^-dependent phosphatidic acid phosphatase lipin 1 (Zhang and Reue, 2017). Lipin 1 transiently associates with the surface of the ER to dephosphorylate phosphatidic acid (PA) to diacylglycerol (DAG), which serves as the precursor for the synthesis of the major membrane phospholipids (phosphatidylcholine or PC and phosphatidylethanolamine or PE) (Figure 1A) (Nohturfft and Zhang, 2009). The regulation of lipin activity and its localization are responsible for controlling the flux of lipids towards ER membrane biogenesis or lipid storage and depend on the proliferative potential of cells (Bahmanyar and Schlieker, 2020; Kwiatek et al., 2020; Zhang and Reue, 2017; Zhang et al., 2014). Thus, lipin is a likely candidate involved in coordinating ER production with mitotic events. There are three lipins in mammalian cells (lipin 1, 2 and 3), and lipin 1 has the highest phosphatidic acid phosphatase (PAP) activity (Grimsey et al., 2008). Lipin 1 is highly regulated by multisite phosphorylation. In mammalian cells, phosphorylation of lipin 1 by the nutrient sensing kinase mTORC1 decreases its affinity for PA and promotes its cytoplasmic enrichment to enable the transcription of genes encoding for fatty acid enzymes by Sterol Regulatory Element Binding Protein-1 (SREBP-1) through an unknown mechanism (Eaton et al., 2013; Peterson et al., 2011). Lipin 1 is also a target of Cdk1, and its hyperphosphorylation in mitosis is consistent with the fact that lipid synthesis is downregulated during the mitotic phase of the cell cycle (Grimsey et al., 2008; Jackowski, 1994). One study in HeLa cells showed a role for lipin 1 in NE breakdown, but this study proposed a local signaling role for DAG production (Mall et al., 2012). Thus, regulation of lipin controls ER membrane biogenesis, but whether control of ER membrane abundance through lipin facilitates mitotic ER reorganization and clearance to prevent chromosomal instability is unknown.

**Figure 1.**
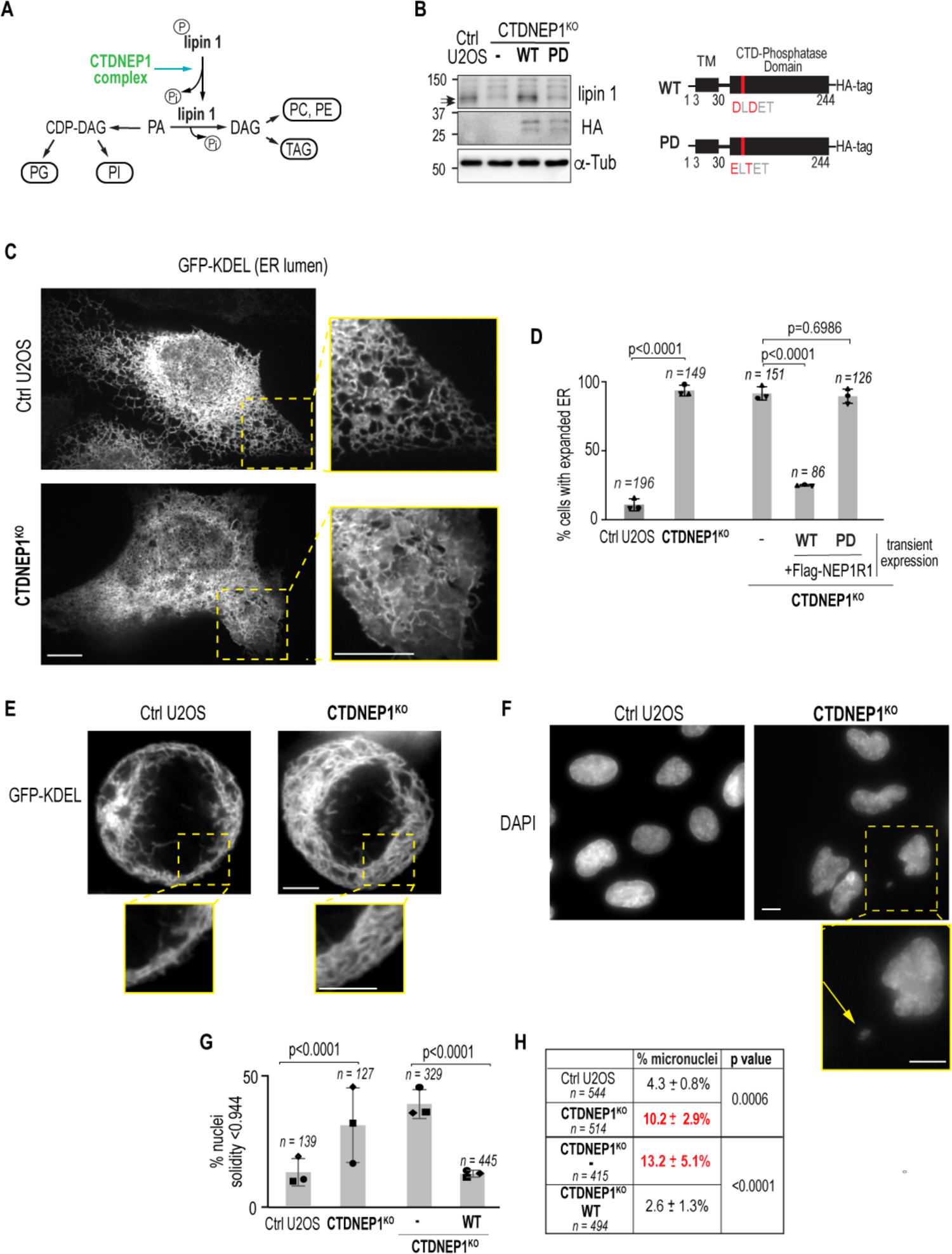
Human CTDNEP1 targets lipin 1 and limits ER membrane expansion and micronuclei formation. **(A)** Schematic showing the function of the CTDNEP1 complex and lipin 1 in lipid synthesis. **(B)** Immunoblot of endogenous lipin 1 from whole cell lysates in indicated cell lines. Schematics (right) show the domain architecture for HA-tagged wild type (WT) and phosphatase dead (PD) CTDNEP1. **(C)** Spinning disk confocal images of GFP-KDEL transiently expressed to mark the ER in indicated cell lines. Scale bars, 10 µm. **(D)** Plot of percentage of cells in indicated cell lines that contain expanded ER membranes as marked by transiently expressed GFP-KDEL. Ctrl and CTDNEP1^KO^ cells were analyzed live. Fixed cell analysis was used for other conditions to confirm expression of indicated epitope tagged CTDNEP1/NEP1R1 constructs. PD = CTDNEP1-HA D67E. n = individual cells, N = 3 experimental repeats for each set of conditions. Mean ± SD shown. P values, Fisher’s exact test of total incidences of expanded ER phenotype. **(E)** Spinning disk confocal images of GFP-KDEL transiently expressed in metaphase cells in indicated cell lines. Mitotic staging was determined by chromatin appearance by DIC (not shown). Scale bars, 10 µm. **(F)** Epifluorescence images of DAPI/Hoescht in indicated cell lines. Arrow marks micronucleus in magnified image. Scale bars, 10µm. **(G)** Plot of incidence of nuclei (n) having a solidity less than 1 standard deviation of control mean solidity value. Mean ± SD shown. The right two and left two bars are quantification of data from separate experiments done on separate days. N = 3 experimental repeats for each set of conditions. P values, Fisher’s exact test of total incidence. **(H)** Incidence of micronuclei in indicated cell lines. The top two and bottom two rows are quantification of data from separate experiments done on separate days. n = number of nuclei, N = 3 experimental repeats. Mean ± SD shown. P values, Fisher’s exact test of total incidence. See also Figure S1 and Figure S2.

To gain insights into whether regulation of ER membrane biogenesis is necessary for mitotic reorganization of the ER to allow accurate chromosome segregation, we focused on the conserved protein phosphatase for lipin 1, known as CTDNEP1 (<u>C</u>-<u>T</u>erminal <u>D</u>omain <u>N</u>uclear <u>E</u>nvelope <u>P</u>hosphatase 1 or Nem1 in fission and budding yeast), because of its role in positively regulating lipin activity in fungi and in *C. elegans* (Bahmanyar et al., 2014; Bahmanyar and Schlieker, 2020; O’Hara et al., 2006; Penfield et al., 2020; Siniossoglou et al., 1998). Little is known about the function of endogenous CTDNEP1 in mammalian cells, although the fact that CTDNEP1 qualifies as a candidate for the long sought-after tumor suppressor in Group 3/4 medulloblastomas (Jones et al., 2012; Northcott et al., 2012) further motivated its functional analysis in human cells. Here, we analyze the function of human CTDNEP1 to demonstrate that the spatial reorganization of ER membranes in mitosis requires limiting ER lipid synthesis to ensure mitotic fidelity. When CTDNEP1 is absent, prometaphase cells fill with aberrantly expanded ER membranes, effectively increasing the viscosity of the mitotic cytoplasm and slowing chromosome movements. The slower mobility of chromosomes during prometaphase corresponds to a reduction in correction of mitotic errors induced by transient spindle disassembly. Mechanistically, we show that CTDNEP1 regulates fatty acid flux towards ER lipid synthesis as a central part of the mTOR-lipin 1 regulatory network. Our understanding of this regulation led to the discovery that treating CTDNEP1 knockout cells with a small molecule inhibitor to the rate limiting enzyme in the *de novo* fatty acid synthesis pathway suppresses ER membrane expansion and micronuclei formation. Together, these data demonstrate that the regulation of ER membrane biogenesis dictates the biophysical properties of mitotic cells to ensure mitotic fidelity. This work further suggests that dysregulated lipid metabolism may be an underlying mechanism that drives aneuploidy in cancer, particularly in the context of cancer-relevant CTDNEP1 truncations and oncogenic mTOR signaling.

## Results

### Aberrantly expanded ER membranes fill mitotic cells and correspond to an increased incidence of micronuclei formation

The conserved integral membrane protein phosphatase CTDNEP1 (yNem1 and *C. elegans* CNEP-1) dephosphorylates lipin (yPah1 and *C. elegans* LPIN-1) in all organisms tested (Bahmanyar et al., 2014; Kim et al., 2007; O’Hara et al., 2006) (Figure 1A); however, almost nothing is known about CTDNEP1’s role in regulation of lipin 1 to control ER membrane biogenesis in mammalian cells. We found that a genome-edited homozygous clonal U2OS knockout cell line for CTDNEP1 as well as cells subjected to RNAi-mediated depletion of CTDNEP1 (Figures S1A-S1B) contain reduced levels of lipin 1 protein as well as the absence of a faster migrating product detected by the endogenous lipin 1 antibody (Figures 1B and S1C). Treatment with exogenous lambda phosphatase resulted in an electrophoretic mobility shift of lipin 1 to a faster migrating species in both wild type and knockout cells (Figure S1D), indicating that lipin 1 is in a mostly phosphorylated form in CTDNEP1 knockout cells. CTDNEP1 belongs to the haloacid dehalogenase (HAD) phosphatase superfamily, whose active sites contain the sequence DXDX(T/V) (Kim et al., 2007; Seifried et al., 2013). Stable expression of an epitope-tagged wild type (WT) CTDNEP1, but not a mutant version of CTDNEP1 with its active site aspartic acid residues mutated to eliminate its phosphatase activity (PD, phosphatase dead), restored both lipin 1 levels and phosphorylation states (Figure 1B). Thus, catalytically active human CTDNEP1 is required to maintain a dephosphorylated pool of lipin 1 and may be required to stabilize lipin 1 protein levels.

Lipid mass spectrometry analysis of CTDNEP1 knockout cells showed increased levels of the major membrane phospholipids (PC and PE) compared to control U2OS cells, and this increase was restored upon stable expression of CTDNEP1-HA (Figure S1E). PC and PE make up the majority of total membrane lipids in cells (van Meer, 2005) and so even small differences in their levels have profound effects on total lipid content. Consistent with this, live imaging of GFP-KDEL revealed an expansion of ER membranes in CTDNEP1 knockout cells in interphase (Figure 1C). In contrast to a mainly tubular ER network in the periphery of control U2OS cells in interphase, membrane sheets and tubules fill the cytoplasm of CTDNEP1 knockout cells (Figure 1C, insets). Expansion of the ER membrane network was also observed with an endogenous ER marker in fixed CTDNEP1 knockout cells (Figure S2A, calnexin) and in U2OS and RPE-1 cells RNAi-depleted for CTDNEP1 (Figures S2B-S2C). Quantitation of segmented ER fluorescence signal confirmed a significant increase of ER membranes in CTDNEP1 knockout cells in interphase (Figure S2B). This phenotype was restored upon transient co-expression of wild type (WT), but not phosphatase defective (PD), CTDNEP1-HA and its binding partner FLAG-NEP1R1 (Figure 1D). Importantly, overexpression of a catalytically active mouse lipin 1β with 19 S/T sites mutated to alanine was sufficient to suppress the expanded ER phenotype resulting from loss of CTDNEP1 (Figure S2D). Furthermore, mitotic CTDNEP1 knockout cells were filled with ER membranes that occupied a greater region of the mitotic cytoplasm than in control cells (Figure 1E). Together, these data demonstrate that the phosphatase activity of human CTDNEP1 acts through catalytically active lipin 1 to limit ER membrane synthesis and further suggests that aberrantly expanded ER membranes crowd the mitotic cytoplasm.

Nuclei in CTDNEP1 knockout U2OS cells appeared less circular than control U2OS cells (Figure 1F). Using solidity (the area fraction of a convex hull for an object) as a parameter for misshapen nuclear morphology, we found a significant percentage of CTDNEP1 knockout cells had low solidity, defined as solidity values less than one standard deviation of the mean solidity value in control U2OS cells (Figure 1G); this phenotype was restored upon stable expression of CTDNEP1-HA (Figure 1G). Thus, there is also a conserved requirement for human CTDNEP1 in maintenance of nuclear structure.

While measuring the nuclear structure defects in CTDNEP1 knockout cells, we frequently observed primary nuclei with micronuclei in interphase (Figure 1F, inset; 10.2 % +/- 2.9 % in CTDNEP1 knockout cells compared to 4.3 % +/- 0.8 % control U2OS cells, Figure 1H) and this phenotype was also restored by stable expression of CTDNEP1-HA (Figure 1H). A major cause of micronuclei formation is the improper attachment of spindle MTs to kinetochores that lead to lagging chromosomes in anaphase (Cimini, 2008; Liu and Pellman, 2020). Thus, the increased incidence of micronuclei in CTDNEP1 knockout cells may be related to excessive ER membranes interfering with some aspect of mitotic chromosomes and their ability to attach to spindle MTs.

### Increased cytoplasmic viscosity in CTDNEP1 knockout cells with excessive ER membranes corresponds to slower chromosome motions in prometaphase

Live imaging of stably expressing GFP-Sec61β and Histone2B-mCherry allowed us to monitor ER occupancy from anaphase onset to mitotic exit in cells RNAi-depleted for CTDNEP1 (Figure 2A; Movie S1). In line with our previous observation (Figure 1E), a consistently greater percentage of the cell diameter was taken up by ER membranes in CTDNEP1 RNAi-depleted cells stably expressing GFP-Sec61β, as well as in CTDNEP1 knockout cells transiently expressing GFP-KDEL (Figures 2A-2B and S3A-S3C), and this corresponded to a smaller region occupied by mitotic chromosomes (Figure 2A-2B). We next imaged synchronized cells transiently expressing GFP-KDEL and Histone2B-mCherry in prometaphase to assess the spatial organization of the excessive ER membranes in relation to unaligned chromosomes (Figure 2C, arrows; Movie S2). In both control U2OS and CTDNEP1 knockout cells in prometaphase, an unaligned chromosome was observed within the peripheral ER network before moving towards the metaphase plate (Figure 2C, arrows; Movie S2); however, unlike control U2OS cells, CTDNEP1 knockout cells contained some ER membranes mislocalized to the spindle region (Movie S2 and Figure 2C). To quantify the incidence of ER membranes invading the spindle region in early phases of mitosis, we enriched for prometaphase and metaphase cells by performing a drugless mitotic shake off directly followed by live imaging of transiently expressed GFP-KDEL to mark the ER and DIC to identify the region occupied by mitotic chromosomes (Figure 2D). Our phenotypic scoring of control and CTDNEP1 knockout cells (“cleared,” “partially cleared,” and “not cleared” in Figure 2D) revealed a significant proportion of CTDNEP1 knockout cells with GFP-KDEL fluorescence signal throughout the mitotic cytoplasm, including the region occupied by mitotic chromosomes as assessed by DIC (Figure 2D). Thus, when CTDNEP1 is absent, excessive ER membranes occupy a larger area of the peripheral mitotic cytoplasm and aberrantly invade the region where mitotic chromosomes are located.

**Figure 2.**
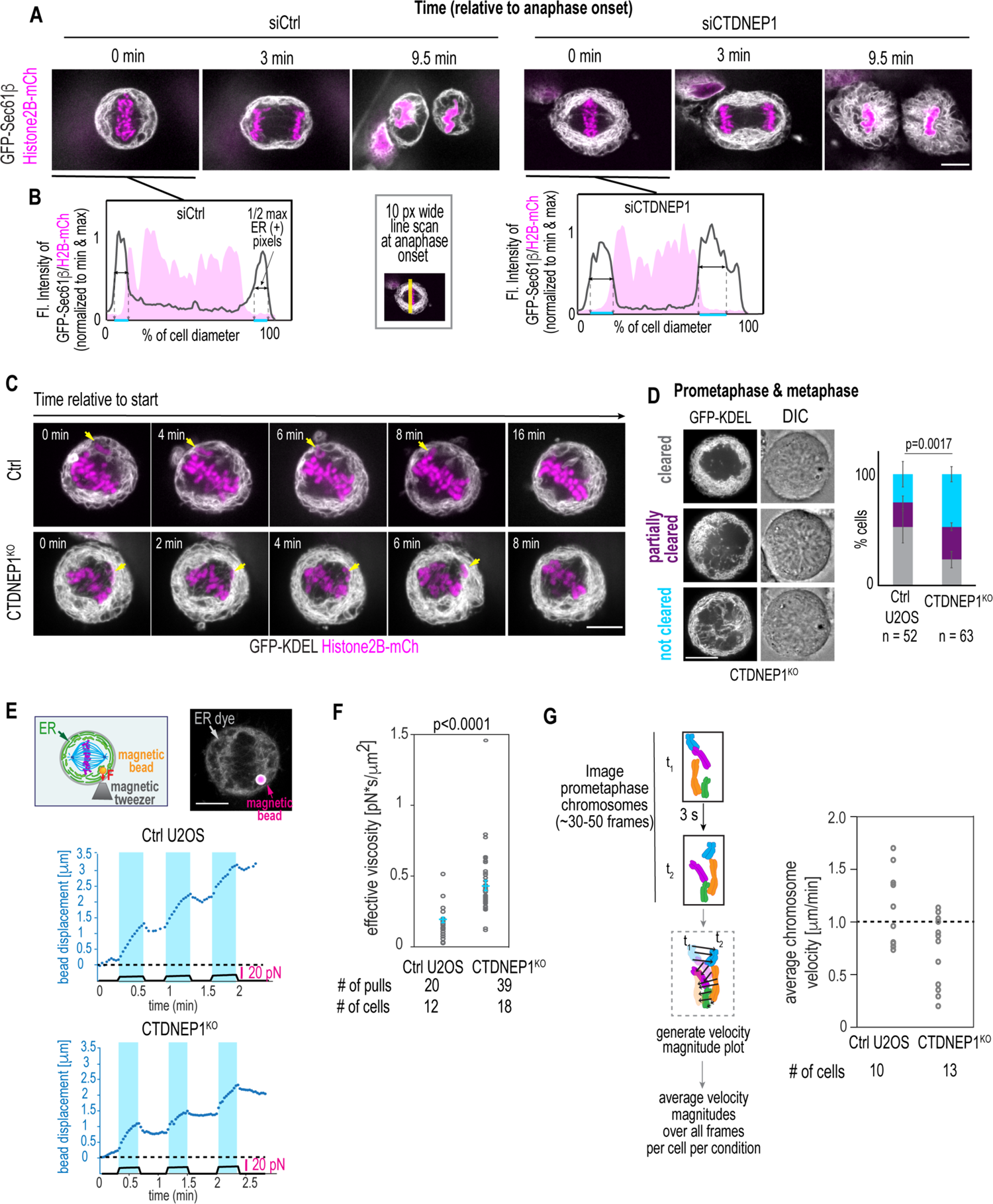
Aberrantly expanded ER membranes fill the mitotic cytoplasm corresponding to an increased effective viscosity and diminished chromosome dynamics. **(A)** Selected spinning disk confocal images of GFP-Sec61β and H2B-mCherry from a time lapse movie of U2OS cells treated with indicated siRNAs. **(B)** Graphs plotting fluorescent intensities of GFP-Sec61β (gray) and H2B-mCherry (magenta) along a 10-pixel line profile drawn along the equatorial region at anaphase onset for indicated conditions. Values are normalized to minimum and maximum values for each channel and to the percentage of the cell diameter. Blue lines indicate percentage of the cell diameter at the half maximum value for GFP-Sec61β. **(C)** Selected spinning disk confocal images from a time lapse movie of GFP-KDEL and H2B-mCh transiently transfected in indicated cell lines after recovery from Cdk1 inhibition. Yellow arrows point to unaligned chromosomes. **(D)** Central plane spinning disk confocal images of transiently expressed GFP-KDEL in CTDNEP1^KO^ U2OS cells with corresponding DIC images (right). Plot: Incidence of indicated phenotypes. Means ± SDs shown. n = cells from N = 3 experimental repeats. P value, χ^2^ test of all phenotype incidences. Scale bars, 10 μm. **(E)** Top: Schematic of experimental set up with magnetic bead and tweezers. Confocal image of ER Tracker Green (gray) and a fluorophore-conjugated magnetic bead (magenta) in a mitotic U2OS cell. Bottom: Plots of applied force (black solid line) and displacement of bead (blue dotted line) in μm. **(F)** Plot of effective cytoplasmic viscosity calculated using force and displacement data collected in E. Individual values (gray circles) and mean ± SEM (blue) are shown. P value, unpaired t test with Welch’s correction. **(G)** Schematic (left) of flow chart to quantify average velocity magnitudes for chromosomes shown for indicated conditions in plot (right). Number of cells measured is shown. Scale bars, 10 μm. See also Figure S3.

We hypothesized that the expanded occupancy of the ER in prometaphase CTDNEP1 knockout cells contributes to viscous forces exerted on mitotic chromosomes. This idea predicts that mitotic chromosomes entrapped by the peripheral ER network would have slower short-range movements, which could impede the proper attachment of dynamic spindle microtubules to kinetochores (Cimini et al., 2003). To test this idea, we monitored the displacement of magnetic beads approximately the size of a mitotic chromosome (∼2-3 μm) in the periphery of mitotic cells in response to a constant force applied with magnetic tweezers (Figure 2E; Movie S3). In control U2OS cells, the displacement of a magnetic bead in response to a constant force of ∼10 pN for ∼20 seconds was ∼1.5 μm (Figure 2E, top plot). In contrast, in CTDNEP1 knockout cells a similar force regime resulted in bead displacement of less than 1 μm (Figure 2E, bottom). In both cases, the relaxation phase after cessation of the applied force was minimal compared to the rising phase, suggesting a greater viscous component influenced bead movement (Figure 2E). These data revealed a two-fold increase in the effective viscosity of the mitotic cytoplasm of CTDNEP1 knockout cells compared to control U2OS cells (Figure 2F; 0.43 ± 0.04 pN * s / μm^2^ in CTDNEP1 knockout compared to 0.19 ± 0.02 pN * s / μm^2^ in control U2OS cells). High temporal resolution imaging of fluorescently-labeled prometaphase chromosomes revealed a greater proportion of CTDNEP1 knockout cells with an average chromosome velocity of less than 1 μm/ min (10/13 CTDNEP1 knockout cells compared to 5/10 control U2OS cells having an average magnitude velocity of <1 μm/ min; Figures 2G and S3D; Movie S4). The reduction in the displacements of mitotic chromosomes in CTDNEP1 knockout cells also suggests an explanation for the more compact occupancy of mitotic chromosomes at anaphase onset (Figure 2B). We conclude that when CTDNEP1 is deleted, excessive ER membranes are more prone to persist in the area occupied by prometaphase chromosomes, effectively increasing the surrounding viscosity to cause slower and overall reduced chromosome motions.

### CTDNEP1 promotes correction of mitotic errors that cause micronucleation

The slow motions of mitotic chromosomes resulting from excessive ER membranes might increase errors in microtubule-chromosome attachments prior to entry into anaphase. This would account for the increased incidence of micronuclei formation we observed in CTDNEP1-deleted cells. Inhibition of the spindle assembly checkpoint (SAC) serves as a readout for the rate of attachment errors because cells enter anaphase without resolving improperly attached or unattached kinetochores (Liu and Pellman, 2020). This results in a substantial increase in the frequency of chromosome missegregation events and micronucleation. To test if loss of CTDNEP1 enhances the frequency of erroneous attachments, we released cells synchronized in late G2 in the presence of an inhibitor to MPS1 (MPSi), the mitotic kinase that activates the spindle assembly checkpoint (Figure 3A, “SAC override”; (Liu et al., 2018)). 39.9 % ± 1.9 % of control cells were micronucleated, reflecting the rate of unresolved errors in attachment prior to entry into anaphase, compared to 43.3 % ± 4.9 % of CTDNEP1 knockout cells were micronucleated (Figures 3B-3C). Although this increase was significant, the difference in the percentage of micronuclei between wild type and knockout cells is small, suggesting that an increased incidence of errors in microtubule attachments to kinetochores may not be the major source of increased micronucleation observed in CTDNEP1-deleted cells.

**Figure 3.**
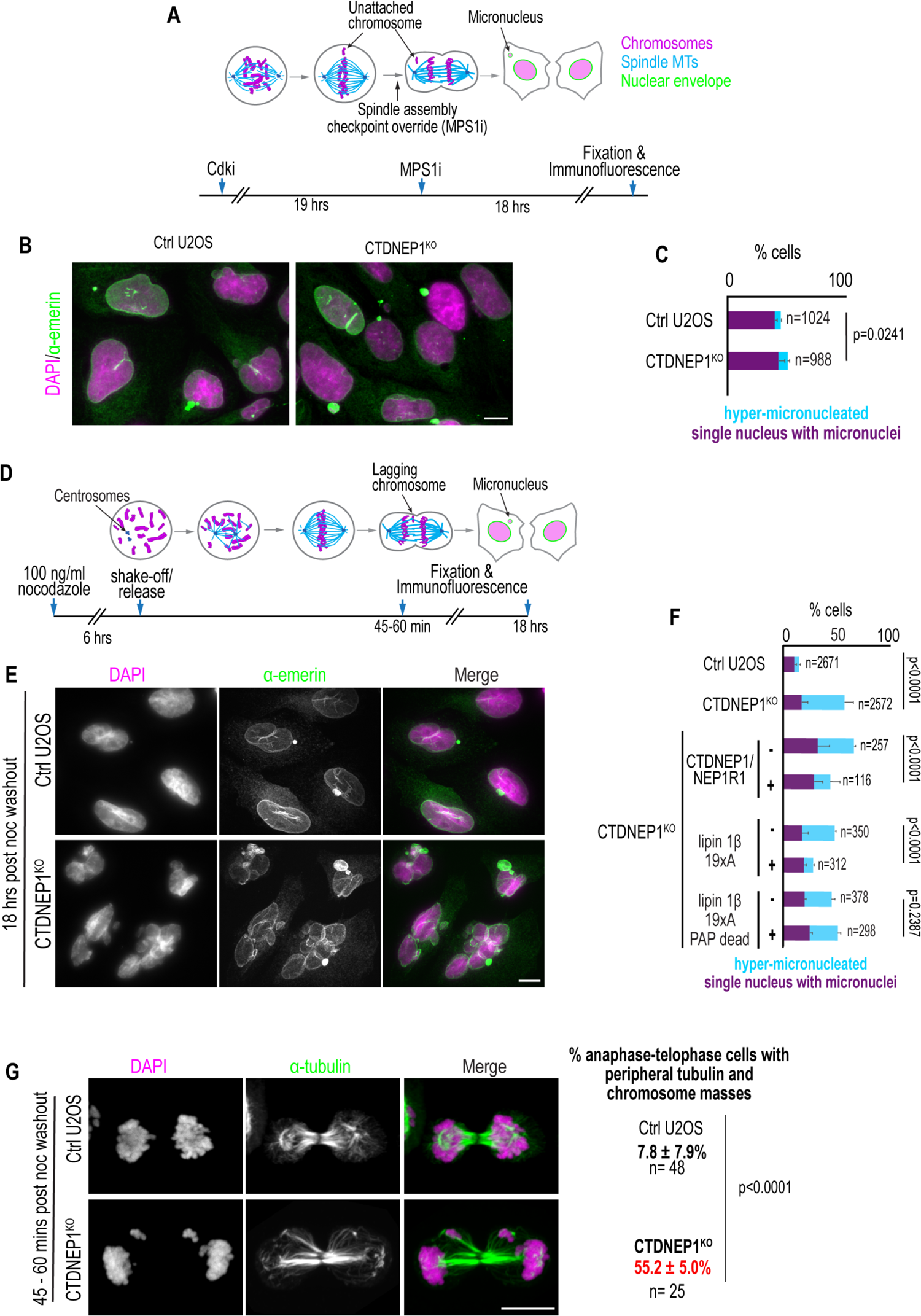
Loss of CTDNEP1 exacerbates the frequency of micronucleation upon transient spindle disassembly. **(A)** Schematic (above) showing spindle assembly checkpoint override and experimental set up (below). **(B)** Maximum projection of confocal images of DAPI/Hoechst (magenta) and anti-emerin (green) in indicated cell lines treated with an inhibitor to MPS1, as indicated in (A). **(C)** Incidence of indicated phenotypes. Mean + SD shown. n = number of nuclei. P value, χ^2^ test of total incidences. **(D)** Schematic (above) of recovery from transient mitotic spindle depolymerization with nocodazole and experimental set up (below) **(E)** Maximum projection of confocal images of DAPI/Hoechst (magenta) and anti-emerin (green) in indicated cell lines after recovery from nocodazole, as indicated in (D). **(F)** Plot of incidences of indicated phenotypes in indicated cell lines and conditions after nocodazole washout for 18 hours as in D. n = number of nuclei. The top two, second two, and bottom four bars are quantification of data from separate experiments done on separate days. Mean + SD shown. P values, χ^2^ test of total incidences. **(G)** Maximum projection of confocal images of control or CTDNEP1^KO^ U2OS cells 45-60 minutes after recovery from nocodazole as in D. Kinetochore-attached microtubules (anti-tubulin, green) and chromosomes (DAPI, magenta) are shown. Anaphase and telophase cells (n) were scored for the indicated phenotypes. Mean ± SD shown. P value, Fisher’s exact test of total incidence. N = 3 experimental repeats. Scale bars, 10 μm. See also Figure S4.

Another possibility is that dampened mitotic chromosome dynamics resulting from an increased viscosity of the cytoplasm in CTDNEP1 knockout cells impedes the correction of errors in MT and kinetochore attachments to cause chromosome missegregation and micronuclei formation. Transient spindle disassembly by washout from nocodazole treatment increases the frequency of merotelic attachments in which a single kinetochore is attached to MTs from both spindle poles instead of just one (Cimini et al., 2003) (Figure 3D). Unlike other mis-attachments, merotelic attachments are not detected by the SAC; however, MT dynamics and chromosomal movements that promote biorientation can substantially reduce these improper attachments prior to anaphase onset resulting in a modest, albeit significant, increase in lagging chromosomes and micronucleation after nocodazole-washout (Cimini, 2008; Cimini et al., 2003) (Figure 3D). Transient spindle disassembly led to the expected increase in micronucleation in control U2OS cells (Figures 3E-3F; compare 10.6 % ± 2.0 % in Figure 3F to 4.3 % ± 0.8 % in Figure 1H; (Cimini et al., 2003)), whereas a significant percentage of CTDNEP1 knockout cells were severely micronucleated - in addition to one or two small micronuclei, these cells contained multiple severely multilobed nuclei (hereafter referred to as “hyper-micronucleation”; 40.0 ± 8.5 % in CTDNEP1 knockout compared to 4.3 ± 1.2 % in control, Figures 3E-3F). The hyper-micronucleation phenotype observed in CTDNEP1 knockout cells as well as RPE-1 cells RNAi-depleted for CTDNEP1 (Figures S4A-S4B) could result from uncorrected merotelic attachments leading to lagging chromosomes in anaphase that remain physically apart from each other upon exit from mitosis. Consistent with this, a significantly greater percentage of CTDNEP1 knockout mitotic cells contained chromosomes that were separated from the main chromosome mass in telophase after recovery from nocodazole treatment (Figure 3G; 7.8 % ± 7.9 % in control U2OS cells compared to 55.2 % ± 5.0 % in CTDNEP1 knockout cells).

**Figure 4.**
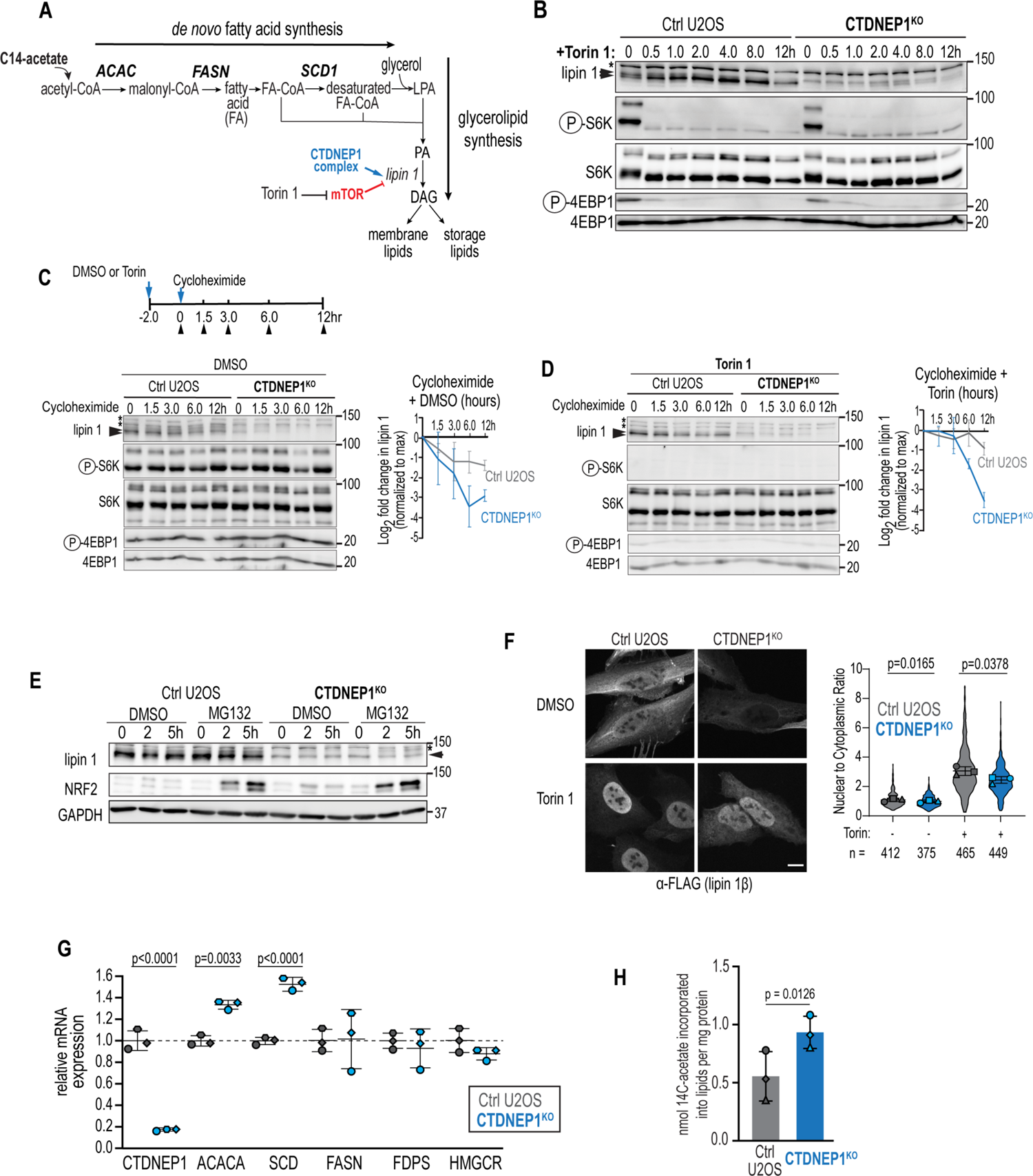
CTDNEP1 controls *de novo* fatty acid synthesis through a mTOR-lipin 1 regulatory network. **(A)** Schematic of steps in *de novo* fatty acid synthesis and glycerolipid synthesis highlighting CTDNEP1 and mTOR regulation of lipin 1. ^14^C-acetate incorporates into acetyl-CoA, and Torin 1 is an ATP-catalytic site inhibitor of mTOR kinase. **(B)** Immunoblot of endogenous lipin 1 from whole cell lysates derived from control and CTDNEP1^KO^ U2OS cells treated with 250 nM Torin 1 for the indicated times. **(C,D)** Immunoblot of whole cell lysates derived from cycloheximide-treated control and CTDNEP1^KO^ U2OS cells pre-treated with DMSO in (C) or 250 nM Torin 1 in (D) for 2 hours and throughout time course. Plots are of fold change in lipin 1 band intensities normalized to the maximum in each condition. Means ± SDs shown. **(E)** Immunoblot of whole cell lysates derived from control and CTDNEP1^KO^ U2OS cells incubated with DMSO or 30 µM MG132 for the indicated times **(F)** Above: Confocal images of anti-FLAG immunostaining in cells expressing FLAG-lipin 1 β treated with DMSO or 250 nM Torin 1 for 4h. Scale bar, 10 μm. Below: Plot of nuclear to cytoplasmic ratio of anti-FLAG fluorescence signal from images taken as in (F). Means ± SDs shown. n = number of cells. P values, paired t test of replicate means. **(G)** qRT-PCR of control and CTDNEP1^KO^ U2OS for genes indicated. Values are normalized to 36B4 expression. Results expressed as fold change in expression relative to mean of control U2OS values. P values, unpaired t tests of ΔCt values. Means ± SD shown. **(H)** Plot of means ± SD of nmol ^14^C-acetate incorporated into lipids per mg of protein in indicated cell lines. P value, paired t test. For all, N = 3 experimental repeats. Arrowheads mark lipin 1 phosphospecies. Asterisks marks non-specific or unchanged bands. See also Figure S5.

The hyper-micronucleation phenotype in CTDNEP1 knockout cells was suppressed upon transient expression of wild type CTDNEP-HA with its binding partner FLAG-NEP1R1 (15.1 ± 9.2% with CTDNEP1/NEP1R1 compared to 34.1 ± 1.6 % with control, Figure 3F). Importantly, overexpression of the 19S/T to A catalytically active, but not phosphatase-dead, lipin 1β, which restores expansion of the ER in CTDNEP1 knockout cells (Figure S2D), also suppressed the exacerbated hyper-micronucleated phenotype (8.7 ± 1.0 % with FLAG-lipin 1β 19xA compared to 30.7 ± 1.4 % in control; 26.2 ± 2.8 % with phosphatase-dead FLAG-lipin 1β 19xA compared to 25.3 ± 3.4 % in control, Figure 3F). Thus, when CTDNEP1-lipin 1-mediated regulation of ER membrane synthesis is absent, enhancing the frequency of merotelic attachments by transient spindle disassembly substantially exacerbates the incidence of lagging chromosomes and formation of micronuclei. Thus, the increased incidence of micronucleation in CTDNEP1 knockout cells results from defects in mitotic error correction. Together, these data suggest that CTDNEP1 restricts ER membrane biogenesis through dephosphorylation of catalytically active lipin 1 in interphase, allowing mitotic cells to inherit a less dense ER network to enable mitotic error correction and prevent micronucleation.

### CTDNEP1 counteracts mTOR-mediated regulation of lipin 1 to limit flux in the *de novo* fatty acid synthesis pathway

To assess the direct link between ER membrane biogenesis and micronucleation, we analyzed the role for CTDNEP1 in regulating lipid synthesis through lipin 1 dephosphorylation. Consistent with previously observed effects of phosphorylation on the membrane associated activity of lipin 1 (Eaton et al., 2013), Mg^2+^-dependent phosphatidic acid phosphatase (PAP) activity was significantly lower in CTDNEP1-deleted cells compared to control U2OS cells (Figure S5A-S5B). Prior work showed that phosphorylation of lipin 1 by mTORC1 positively regulates *de novo* fatty acid synthesis through SREBP-1-mediated transcription of fatty acid synthesis enzymes (Peterson et al., 2011). We reasoned that CTDNEP1 may dephosphorylate lipin 1 to counteract this pathway and limit fatty acid synthesis (Figure 4A). Our data demonstrating that expression of lipin 1β with 19 S/T phosphorylation sites mutated to alanine, which includes the sites targeted by mTORC1 (Peterson et al., 2011), is sufficient to restore the ER expansion (Figure S2D) and hyper-micronucleation (Figure 3F) phenotypes of CTDNEP1 knockout cells further supports the possibility that CTDNEP1 functions through the mTOR-lipin 1 regulatory network to limit ER membrane biogenesis and ensure mitotic fidelity. We thus monitored the phosphorylation state of lipin 1 in a time course of control and CTDNEP1 knockout cells treated with the mTOR kinase inhibitor Torin 1 (Peterson et al., 2011). Control U2OS cells showed a doublet of endogenous lipin 1 representing its distinct phosphorylation states (Figure 4B). Treating U2OS cells with Torin 1 for 30 minutes and after up to 12 hours caused a shift in the electrophoretic mobility of lipin 1 to the faster migrating, dephosphorylated product consistent with rapid dephosphorylation of lipin 1 in the absence of mTOR kinase activity (Figure 4B). In contrast, the dephosphorylated species of lipin 1 did not appear until 2 hours of Torin 1 treatment in CTDNEP1 knockout cells and was present at much lower levels throughout the time course of the experiment when compared to control U2OS cells (Figure 4B). Thus, the majority of lipin 1 that becomes dephosphorylated upon inhibition of mTOR kinase activity depends on CTDNEP1, indicating that CTDNEP1 is the major phosphatase counteracting mTOR phosphorylation of lipin 1 and supporting the idea that CTDNEP1 restricts ER membrane expansion as a part of the mTOR-lipin 1 regulatory network to ensure mitotic fidelity.

mTOR phosphorylation of exogenously expressed lipin 1 prevents its nuclear enrichment to allow SREBP-1-target gene expression (Peterson et al., 2011) and primes it for subsequent targeting to the proteasome for degradation (Shimizu et al., 2017). We found that CTDNEP1 dephosphorylation of endogenous lipin 1 stabilizes lipin 1 protein levels by preventing its proteasomal degradation and promoting its nuclear translocation. Cycloheximide treatment to prevent new protein translation showed that lipin 1 protein levels are less stable in CTDNEP1 knockout cells in both untreated (Figure 4C) and Torin 1-treated conditions (Figure 4D). Small molecule inhibition of the proteasome with MG132 results in the accumulation of a slower-migrating lipin 1 product in CTDNEP1 knockout cells that is likely a hyperphosphorylated form rapidly degraded under normal conditions (Figure 4E). Thus, CTDNEP1 prevents targeting of a pool of lipin 1 for degradation by the proteasome. CTDNEP1 knockout cells also contained lower levels of nuclear FLAG-lipin 1β under normal conditions as well as after 4 h of Torin 1 treatment (Figure 4F). Furthermore, the SREBP-1-target genes encoding for acetyl-CoA carboxylase alpha (ACACA), which functions in the first committed step in *de novo* fatty acid synthesis by catalyzing the carboxylation of acetyl-CoA to malonyl-CoA, as well as Stearoyl-CoA desaturase (SCD), the rate limiting enzyme in fatty acid desaturation, are increased in CTDNEP1 knockout cells (Figure 4A and 4G), although the levels of the SREBP-1-target gene Fatty Acid Synthase (FASN) remained unchanged under these steady state conditions (Figure 4G) (Nohturfft and Zhang, 2009). Consistent with prior data showing that regulation of SREBP2 targets is not mediated by lipin 1 (Peterson et al., 2011), (FDPS (farnesyl diphosphate synthase) and HMGCR (3-hydroxy-3-methylglutaryl-coenzyme A reductase) were not detectably different in CTDNEP1 knockout and control cells (Figure 4G) (Lamming and Sabatini, 2013). These data suggest that CTDNEP1 knockout cells undergo increased *de novo* fatty acid synthesis. To directly test this idea, we monitored the incorporation of radiolabeled acetate in the lipid fraction after 5 hours of incubation (Figure 4H). Lipids extracted from CTDNEP1 knockout cells contained nearly two-fold more radiolabel than control U2OS cells, while background acetate labeling was similar between control and CTDNEP1 knockout cells (Figures 4H and S5C). Together, these data reveal that control of lipin 1 stability and localization by a CTDNEP1-lipin 1-mTOR regulatory network maintains levels of fatty acid synthesis at least in part through transcription of the rate limiting enzymes in the *de novo* fatty acid synthesis pathway and thus might be required to coordinate ER lipid synthesis with mitotic reorganization of the ER and mitotic fidelity.

### Mitotic errors in CTDNEP1 knockout cells result from excessive ER membrane biogenesis caused by increased flux in the *de novo* fatty acid synthesis pathway

Our mechanistic analysis suggested that targeting the rate-limiting step of the *de novo* fatty acid synthesis pathway may directly assess the link between expanded ER membranes and micronucleation phenotypes of CTDNEP1 knockout cells. We first determined if the expanded ER membrane phenotype in CTDNEP1 knockout cells indeed results from an increase in *de novo* fatty acid synthesis. Treating cells with a small molecule inhibitor that targets acetyl-CoA carboxylase (ACAC) (TOFA, 5’(Tetradecyloxy)-2-furoic acid) (McCune and Harris, 1979) abolished *de novo* fatty acid synthesis within 5 hours (Figures 5A and S5C) and led to a substantial decrease in ER network density in control U2OS cells after 24 h (Figures 5B-5C and S6A-S6B). Rather than the relatively even distribution of ER tubules throughout the cell periphery in untreated cells (“normal” in Figures 5B-5C and S6A), the remaining ER membranes were mainly concentrated near the NE in control cells treated with TOFA (“reduced” in Figures 5B-5C and S6A). The addition of exogenous fatty acids (1:2:1 palmitic:oleic:lineoleic acids conjugated to fatty acid-free BSA) did not alter the morphology of the ER on its own (Figure S6A) but suppressed the altered ER morphology in control U2OS cells treated with TOFA, indicating that the “reduced” ER phenotype resulted from a reduction in the flux of fatty acids towards the synthesis of ER membrane lipids (Figures 5A-5C and S6A, S6C). TOFA treatment of CTDNEP1 knockout cells caused a wild type appearance of the ER network that expanded with the addition of exogenous fatty acids (Figures 5B-5C and S6A-S6B). In addition to suppressing the expanded ER phenotype in CTDNEP1 knockout cells, TOFA treatment suppressed abnormal nuclear structure (Figure 5D). These results indicate that the expanded ER phenotype and altered nuclear morphology in CTDNEP1 knockout cells results from excessive incorporation of fatty acids into ER membrane lipids and provides us with a tool to directly determine if suppressing ER membrane biogenesis is sufficient to prevent mitotic defects.

**Figure 5.**
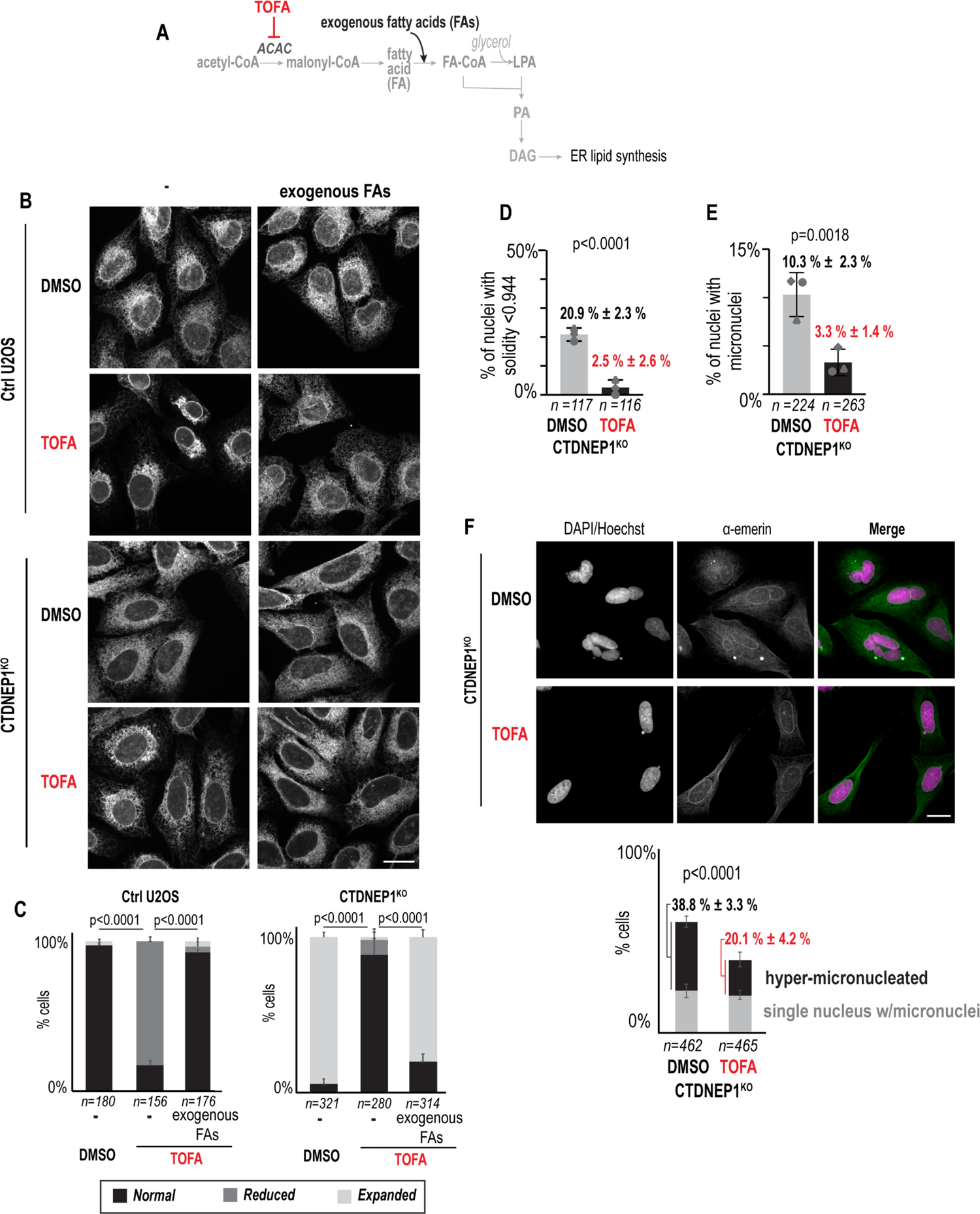
Small molecule inhibition of ACAC suppresses the expansion of the ER and micronuclei formation resulting from loss of CTDNEP1. **(A)** Schematic highlighting the target of the small molecule inhibitor TOFA on the rate-limiting enzyme that functions in the production of malonyl-CoA for *de novo* fatty acid synthesis. The step in *de novo* fatty acid synthesis where exogenous fatty acids are incorporated downstream of ACAC is shown. **(B)** Maximum projections of confocal images of anti-calnexin in fixed and immunostained indicated cell lines treated with DMSO or 10 µM TOFA for 24 hours with and without exogenous fatty acids. Scale bar, 20µm. **(C)** Plot of incidences of anti-calnexin immunostaining in cells showing a wild type ER appearance (“Normal,” black), abnormally reduced ER appearance (“reduced”, dark gray), or expanded ER appearance (“expanded,” light gray). n = number of cells. Means ± SDs shown. P values, χ^2^ test of total incidences. **(D)** Incidence of cells (n) with micronuclei after 48 hours of 10 µM TOFA or DMSO treatment. Means ± SDs shown. P values, Fisher’s exact test. **(E)** Incidence of nuclei (n) with solidity less than one SD from mean solidity value of control U2OS cells after 24 hours of treatment with 10 µM TOFA or DMSO. Means ± SDs shown. P values, Fisher’s exact test. **(F)** Above: Confocal images of DAPI and anti-emerin immunostaining of fixed CTDNEP1^KO^ cells treated with DMSO or 10 μM TOFA for 24 hours during nocodazole washout as in Figure 3D-3F. Scale bar, 20µm. Plot: Incidence of specified phenotypes. n = number of cells. Means and SDs are shown. P value, χ^2^ test of indicated phenotypes. For all panels, N = 3 experimental repeats. See also Figure S6.

Importantly, in addition to ER morphology and nuclear structure, small molecule inhibition of ACAC in CTDNEP1 knockout cells suppressed the increased incidence of micronuclei (Figure 5E, 10.3 % ± 2.3 % in DMSO-treated cells compared to 3.3% ± 1.4% in TOFA-treated cells) as well as the hyper-micronucleation phenotype that occurs after transient spindle disassembly (Figure 5F, 38.8% ± 3.3 % in DMSO-treated versus 20.1% ± 4.2 % in TOFA-treated cells). Thus, we conclude that limiting the production of ER membranes through downregulation of *de novo* fatty acid synthesis downstream of mTOR-mediated signaling is necessary for mitotic fidelity.

## Discussion

Here, we show restricting ER membrane biogenesis is necessary for correction of mitotic errors and prevention of micronucleation. Specifically, restricting ER lipid synthesis lowers the viscosity of the mitotic cytoplasm to enable chromosome motions necessary for biorientation (Figure 6). Mechanistically, we show that CTDNEP1 restricts ER membrane biogenesis by counteracting mTOR to stabilize the levels of the phosphatidic acid phosphatase lipin 1 and limit *de novo* fatty acid synthesis. We demonstrate that increased flux of *de novo* fatty acid synthesis expands the ER and limits mitotic error correction (Figure 6). Together, our findings reveal a novel aspect of ER regulation necessary for mitotic fidelity and expand our understanding of how the spatial organization of the ER in mitotic cells controls mitotic processes. Our work further suggests a potential link between cancer-associated truncations in CTDNEP1 and oncogenic mTORC1 signaling to aneuploidy through dysregulation of lipid metabolism.

**Figure 6:**
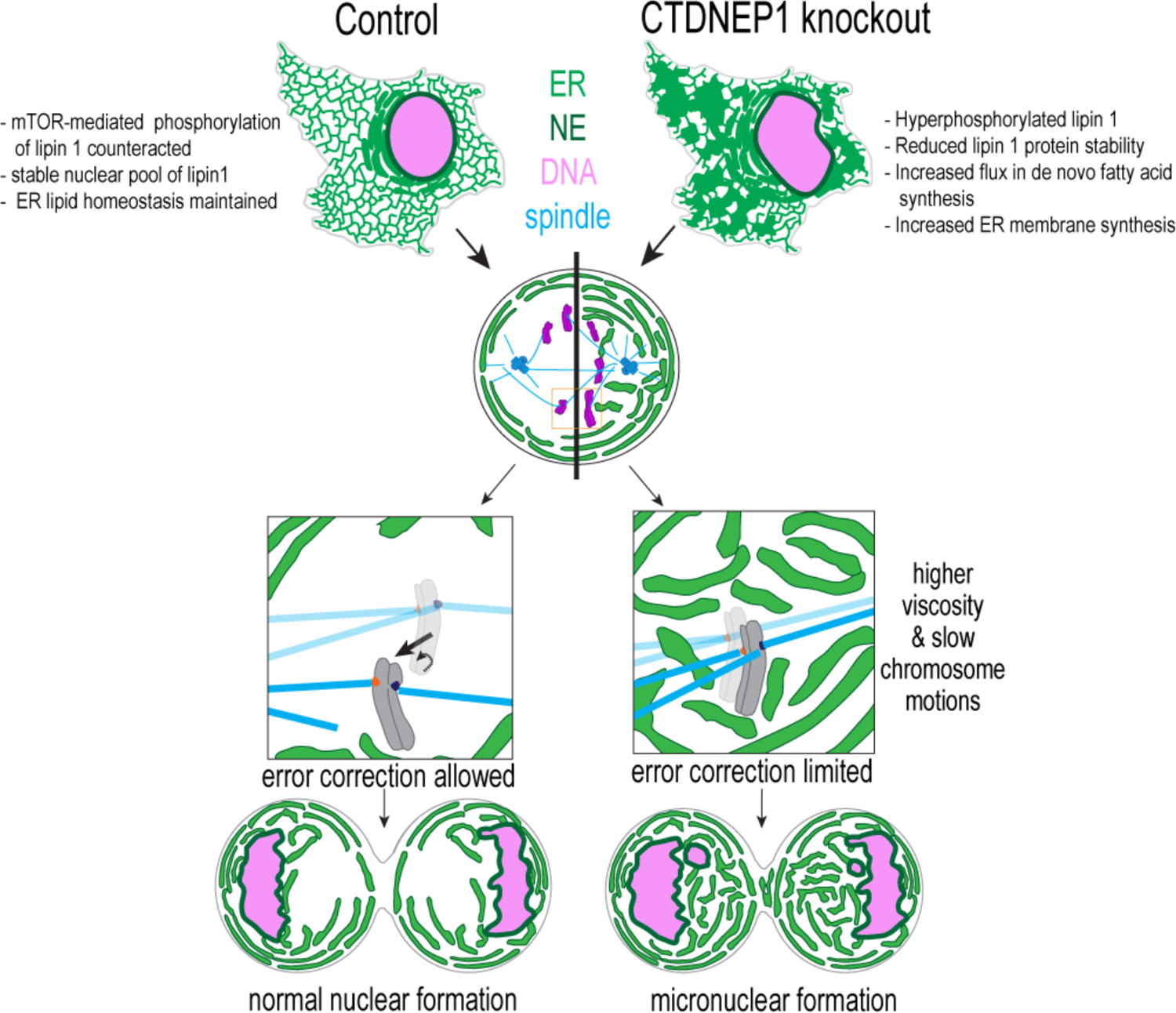
Restricting ER membrane biogenesis through a CTDNEP1-lipin 1-mTOR regulatory network enables chromosome movements necessary for biorientation. Above: When CTDNEP1 is absent, mTOR phosphorylation of lipin 1 prevails and leads to decreased lipin 1 stability, increased flux in *de novo* fatty acid synthesis and increased ER membrane biogenesis. Excess ER membranes in mitosis contribute to higher cytoplasmic viscosity and dampened chromosome motions. The lack of chromosome motions limits mitotic error correction, leading to micronuclei formation (below).

Our data indicate that ER membrane reorganization dictates the biophysical properties of mitotic cells essential to animal cells in which the cytoplasm and membrane-bound organelles are shared by mitotic chromosomes after NE breakdown (Liu and Pellman, 2020). Fungi undergo a closed mitosis and so expansion of the NE, which occurs through phospho-regulation of Pah1/Ned1, accommodates the elongating mitotic spindle that forms inside the nucleus to segregate chromosomes (Makarova et al., 2016; Webster et al., 2009). The rapid divisions of the early *C. elegans* embryos results in penetration of the partially disassembled NE by mitotic spindle MTs; ER membranes in *cnep-1* mutants wrap the NE to delay NEBD but do not interfere with the chromosome segregation machinery (Bahmanyar et al., 2014). Thus, the coordination of ER membrane biogenesis with mitotic reorganization of the ER to promote accurate chromosome segregation is an essential feature of animal cells that have evolved to undergo open mitosis.

In open mitosis, restricting the abundance and the occupancy of the ER within the periphery of mitotic cells maintains a lower viscosity of the mitotic cytoplasm to enable chromosome movements necessary for mitotic error correction (Figure 6). Our data showing that the frequency of lagging chromosomes and micronuclei formation upon transient spindle disassembly is substantially increased under conditions when ER membranes are expanded suggests that the dampened chromosome dynamics resulting from the increase in the effective viscosity of the mitotic cytoplasm impedes the correction of merotelic orientations (Figure 6) (Cimini et al., 2003). Merotelic attachments go unnoticed by the spindle assembly checkpoints (Cimini, 2008) and so they present a unique threat to rapidly dividing cells and are a major source of aneuploidy in cancer cells (Cimini, 2008; Cimini et al., 2002; Cimini et al., 2001). In a merotelically-oriented kinetochore, loss of attachment from one pole can allow the chromosome to biorient - forces from MTs on the attached kinetochore promote the chromosome to move so that the unattached kinetochore faces the proper spindle pole (Cimini, 2008; Cimini et al., 2003) (Figure 6). We propose that slower chromosome mobilities resulting from the increase in viscosity of the surrounding thickened ER network impedes error correction by slowing chromosome biorientation, resulting in re-attachment of the kinetochore to the incorrect pole (Figure 6). Our magnetic tweezer experiments further support this idea by showing that the application of a constant force on a magnetic bead that is approximately the size of a mitotic chromosome and is ensheathed in the thick ER network results in reduced bead displacement in CTDNEP1 knockout cells. The magnitude of the applied forces with magnetic tweezers are relevant to biological force scales, allowing us to infer the influence of the surrounding ER on movements of chromosomes in response to forces that may be similar to spindle MT pulling forces on a kinetochore (Rago and Cheeseman, 2013). CTDNEP1-deleted cells exit mitosis despite improper MT attachments, leading to a “hyper-micronucleated” phenotype, possibly because merotelic attachments are undetected by spindle checkpoints and/or because prolonged mitotic delays can cause “mitotic slippage” (Cimini, 2008; Rieder and Maiato, 2004).

To understand how ER membrane biogenesis is linked to mitotic fidelity, we show for the first time that human CTDNEP1 is the major phosphatase that counteracts mTOR phosphorylation of lipin 1. Prior work *in vitro* suggested that CTDNEP1 dephosphorylates a site on lipin 1 later confirmed to be targeted by mTOR (Wu et al., 2011). Work in budding yeast showed the dephosphorylation of Pah1 upon inhibition of Tor requires the presence of the Nem1/Spo7 complex (Dubots et al., 2014). We demonstrate that human CTDNEP1 antagonizes mTOR-mediated regulation of lipin 1 to stabilize a nuclear pool of lipin 1 and limit the transcription of rate-limiting enzymes in the *de novo* fatty acid synthesis pathway (ACACA and SCD1). We suggest that this regulation is critical to coordinating ER membrane synthesis in highly proliferative cells to ensure mitotic fidelity. Whether the hyperphosphorylated lipin in CTDNEP1-deleted cells that retains some lipin PAP activity together with the increase we observe in ACACA transcript levels is sufficient to have the profound effect on fatty acid synthesis and ER membrane expansion is unclear. The fact that nearly complete inhibition of *de novo* fatty acid synthesis for 24 h causes a severe reduction in ER membranes in control cells, but causes the ER of CTDNEP1-deleted cells to have a more normal appearance, suggests that a slower rate of lipid breakdown may also be involved in this regulation. Future work is required to understand the feedback regulation of the CTDNEP1-lipin 1-mTOR regulatory network that coordinates ER membrane biogenesis with mitotic ER reorganization to prevent mitotic errors.

Our findings that loss of human CTDNEP1 contributes to chromosome instability through dysregulation of lipid metabolism is highly relevant to the fact that Group 3/4 medulloblastomas with high levels of genome instability frequently carry truncation mutations in CTDNEP1 (Jones et al., 2012; Northcott et al., 2012). This subtype contains a wild type copy of *TP53* but is characterized by frequent loss of heterozygosity of 17p (where both *TP53* and *CTDNEP1* reside), and so the identity of a candidate tumor suppressor in this region of 17p has been long sought after (Jones et al., 2012; Northcott et al., 2012). Our data showing that loss of CTDNEP1 causes chromosomal instability and formation of micronuclei suggests that CTDNEP1 may function as a tumor suppressor and when mutated may contribute to the malignancy of this cancer. Micronuclei occur frequently in cancer, and their membranes are prone to rupture, causing cancer-relevant chromosome rearrangements and activation of proinflammatory pathways (Hatch et al., 2013; Liu and Pellman, 2020; Ly and Cleveland, 2017; Mackenzie et al., 2017; Zhang et al., 2013). Increased *de novo* fatty acid synthesis is a hallmark of many cancers; however, why this is an advantage to tumor cells is not fully understood (Currie et al., 2013). Our data links increased fatty acid synthesis to micronuclei formation and defines the mechanistic relationship between these processes. This work additionally suggests that aneuploidy in the context of oncogenic mTOR signaling may be mediated by misregulated lipid biosynthesis (Lamming and Sabatini, 2013). Together, these findings suggest that the use of inhibitors of enzymes in the *de novo* lipid synthesis pathway may be a potential therapeutic strategy in cancers with chromosomal instability. Uncovering the significance of CTDNEP1 and mTOR-mediated upregulation of ER lipid synthesis in the context of chromosomal instability in cancer will be an exciting and important avenue of future research.

## Acknowledgements

We thank M. Deline for initial experiments that guided experiments; M. Deline, L.K. Schroeder, and C. Hu for cloning of some constructs used in the study; S. Lee for image analysis assistance; G. Celma for efforts toward a CTDNEP1 antibody, P. Forscher for the use of a custom heated stage insert for some live cell imaging experiments, and I. Cheeseman for helpful discussion. This work was supported by an NIH R01 (GM131004) and NSF CAREER Award (1846010) to S. Bahmanyar, an NSF MRI (DBI-1919834) to D. Needleman, and NIH R01 (GM136900) to T. Harris. Additional support is to H. Merta by NIH (T32 GM100884 and T32 GM007499) and the Gruber Foundation, to J.W. Carrasquillo Rodríguez by NIH (T32 GM722345), to M. Anjur-Dietrich by an NSF Quantitative Biology Student Fellowship (1764269), and to M. Granade by an American Heart Association Fellowship (20PRE35210847).

## Author contributions

H. M generated the CTDNEP1 knockout cell line and fluorescent stable cell lines and performed most imaging experiments and lipidomic data analysis, J.W.C.R generated stable cell lines and performed experiments related to mTOR, T.V performed experiments related to lipin localization, M.I.A-D (supervised by D.N) performed the magnetic tweezer experiments and chromosome mobility imaging and analysis, M.E.G (supervised by T.E.H) performed the radiolabeling, lipid extraction and PAP activity experiments. H.M and S.B wrote the manuscript. S.B supervised the project.

## Supplemental Figures and Figure Legends

**Figure S1.**
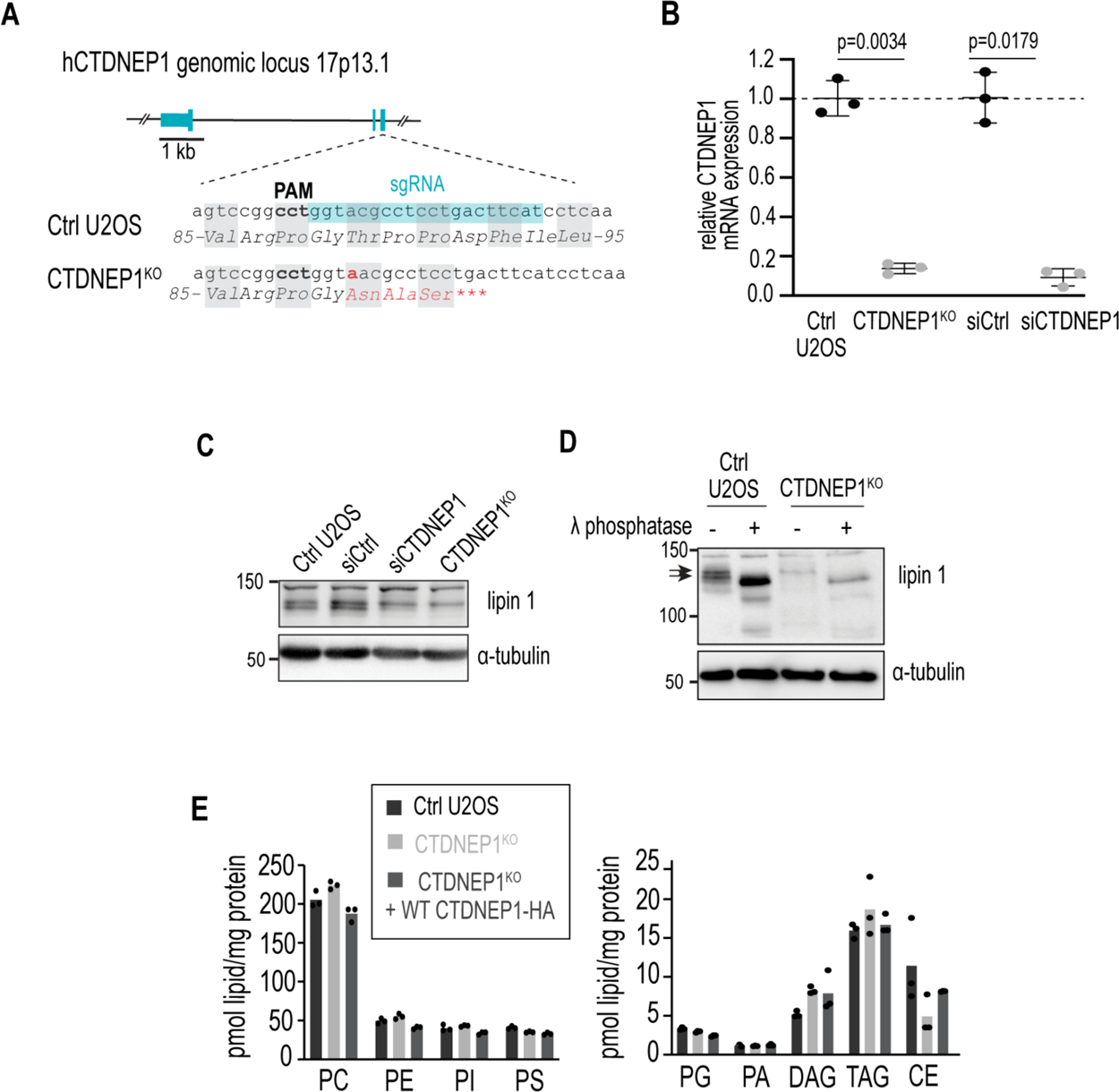
Human CTDNEP1 restricts ER membrane biogenesis through regulating lipin 1 phosphorylation and activity, pertaining to figure 1. **(A)** Schematic showing CRISPR-Cas9 strategy for knocking out endogenous human CTDNEP1 in U2OS cells. sgRNA targeted region (blue) and genome-edited DNA and corresponding mutated amino acid sequence (red) are shown. PAM = protospacer adjacent motif. **(B)** qRT-PCR of CTDNEP1 expression in U2OS cell lines with indicated treatments. siRNA-treated cells are from the same experiment as Figure S2B-S2C. Values are normalized to GAPDH expression. Results expressed as fold change in expression relative to mean of control U2OS or siCtrl-treated U2OS values. P values, paired t tests of ΔCt values. Means ± SD shown. **(C)** Immunoblot of endogenous lipin 1 from whole cell lysates derived from control or CTDNEP1^KO^ U2OS and control U2OS treated with indicated siRNA. **(D)** Immunoblot of lipin 1 in whole cell lysates of indicated cell lines treated with lambda phosphatase. Arrows point to lipin phosphospecies bands collapsed with phosphatase treatment. **(E)** Plots of pmol of lipids determined by mass spectrometry lipid profiling normalized to protein concentration from 3 technical repeats per condition.

**Figure S2.**
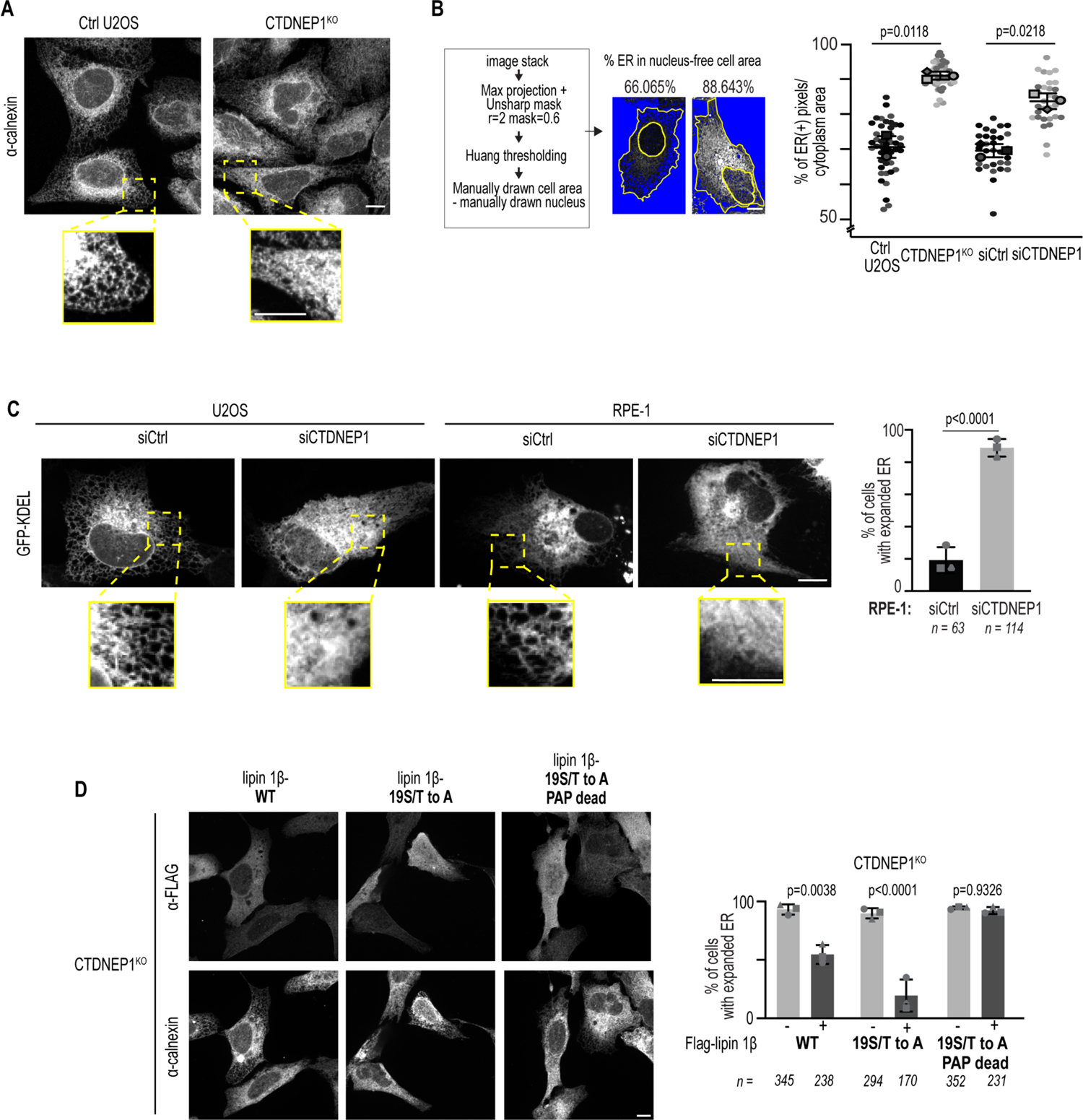
CTDNEP1 restricts ER membrane biogenesis through regulating lipin 1 phosphorylation and activity, pertaining to figure 1. **(A)** Maximum projection of confocal images of calnexin staining in control and CTDNEP1^KO^ cells. Inset brightnesses adjusted to lower threshold than uncropped image to highlight fine ER morphology. **(B)** Schematic: Images of GFP-KDEL in transiently expressing cells were processed by unmask sharpening and thresholded, and cell/nuclear borders were drawn manually. Plot: Graph of percent area of KDEL-positive pixels in the cell area - nuclear area for cells (n) from indicated conditions. Individual data points and means ± SD shown. P values, paired t tests of replicate means. **(C)** Left: Maximum projections of confocal images of transiently-expressed GFP-KDEL in U2OS or RPE-1 cells treated with indicated siRNA. Inset brightnesses adjusted to lower threshold than uncropped image to highlight fine ER morphology. Right: Plot of incidence of expanded ER in cells treated with indicated siRNAs. Means ± SD shown. n = number of cells. P value, Fisher’s exact test of total incidences. **(D)** Confocal images of calnexin and FLAG staining in U2OS CTDNEP1^KO^ cells transfected with the indicated lipin 1β vectors and GFP-KDEL as a co-transfection marker (not shown). Plot: Incidence of expanded ER (calnexin). n = number of cells. Means ± SD shown. P values, Fisher’s exact test of total incidences. For all images, scale bars are 10 μm. For all quantification, N = 3 experimental repeats.

**Figure S3.**
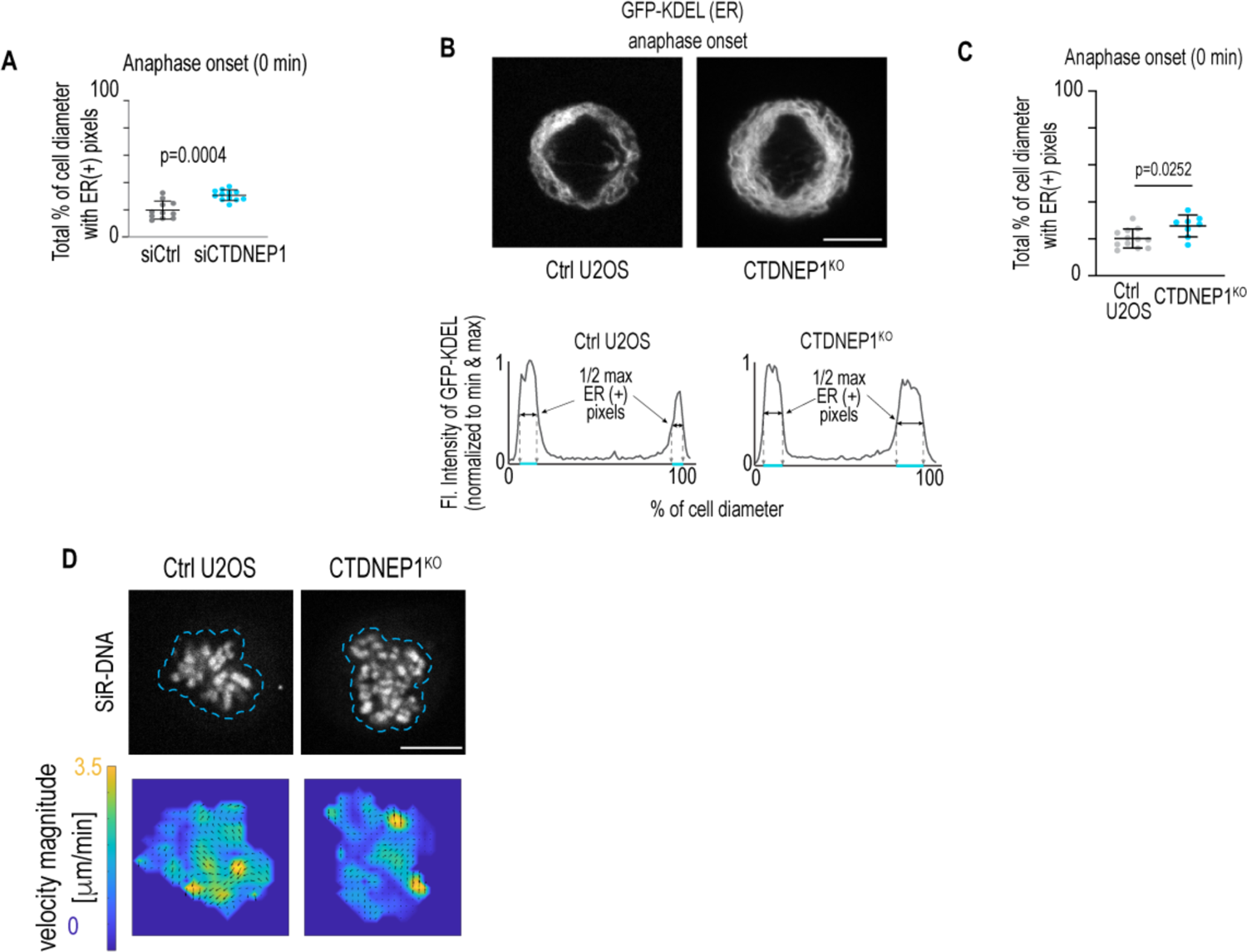
Ectopic ER membranes fill mitotic cytoplasm in CTDNEP1 knockout U2OS cells, pertaining to Figure 2. **(A)** Plot of percent of cell equatorial diameter occupied by Sec61β (gray)- positive pixels from images as in Figure 2A. Images were collected from 7 experiments. Individual values (per cell), means ± SDs shown. P value, Mann-Whitney 2-tailed unpaired t test. **(B)** Images: Confocal images from time lapse movies of GFP-KDEL in transiently-expressing cells. Cells are determined to be at anaphase onset by chromatin appearance using SiR-DNA staining (not shown). Plots: Graphs plotting fluorescent intensities of GFP-KDEL along a 10-pixel line profile drawn along the equatorial region at anaphase onset for indicated conditions. Values are normalized to minimum and maximum values for each channel and to the percentage of the cell diameter. Blue lines indicate percentage of the cell diameter at the half maximum value for GFP-KDEL. **(C)** Plot of percent of cell diameters occupied by GFP-KDEL-positive pixels Individual values (per cell), means ± SDs shown. N = 3 experimental repeats. P value, Mann-Whitney 2- tailed unpaired t test. **(D)** Image: confocal images of SiR-DNA staining in cells in prometaphase for measurement of chromosome velocity. Blue outline shows area shown in the corresponding velocity magnitude fields (below) for frames including top images and subsequent frame (3 s later). For all images, scale bars are 10 μm.

**Figure S4.**
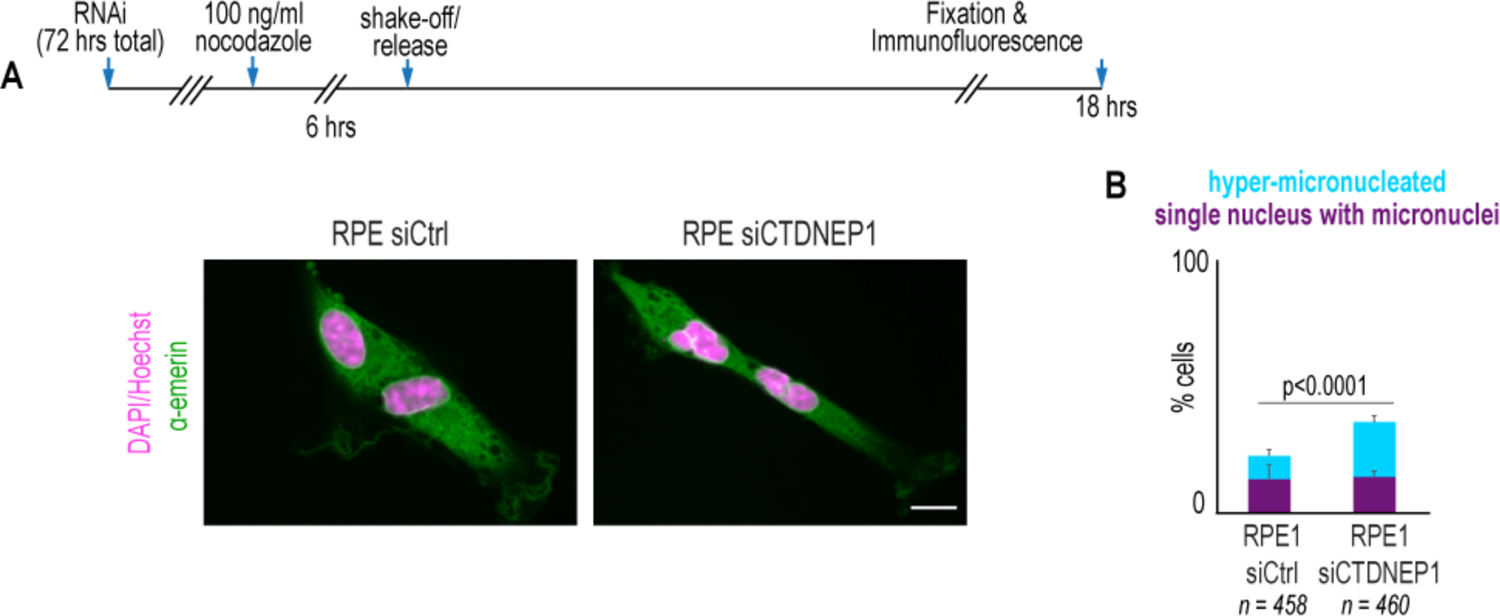
CTDNEP1 limits mitotic errors in response to recovery from spindle depolymerization in RPE-1 cells, pertaining to Figure 3. **(A)** Confocal image of emerin and DAPI/Hoechst staining in RPE-1 cells treated as shown. Scale bar, 10 µm. **(B)** Incidence of indicated phenotypes. Means + SD shown. n = number of cells from N = 3 experimental repeats. P value, Chi squared test of total incidences of indicated phenotypes.

**Figure S5.**
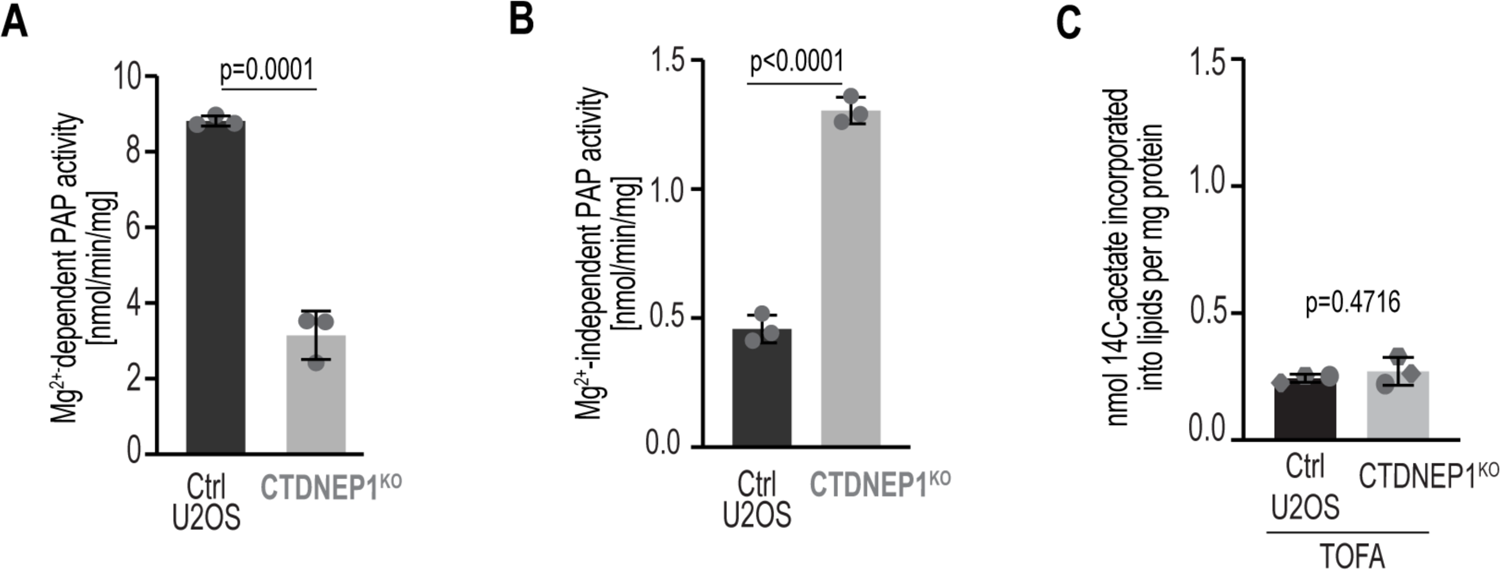
Cellular PAP activity and basal acetate incorporation in control and CTDNEP1^KO^ cells, pertaining to Figure 4. **(A)** Plot of Mg^2+^-dependent phosphatidic acid phosphatase activity in indicated cell lines. Means ± SD shown. P value, unpaired t test. n = 3 samples per condition. **(B)** Plot of Mg^2+^-independent phosphatidic acid phosphatase activity in indicated cell lines. n = 3 samples per condition. Means ± SD shown. P value, unpaired t test. **(C)** Plot of acetate incorporation into lipids of cells treated with 10 µM TOFA. Taken from same dataset and used to calculate background acetate incorporation in Figure 4H. Means ± SD shown. N = 3 experimental repeats. P value, paired t test.

**Figure S6.**
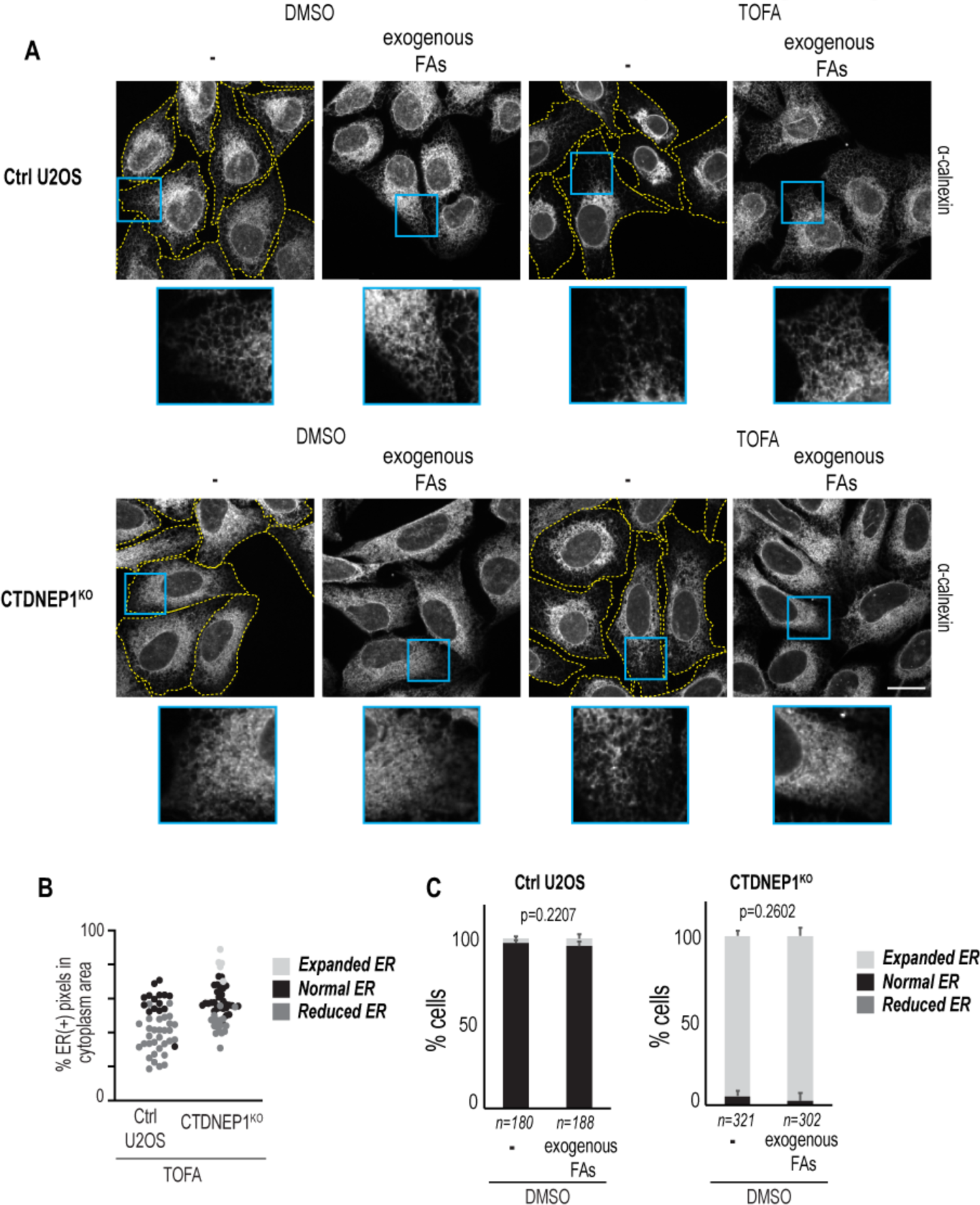
Analysis of ER phenotypes upon inhibition of *de novo* fatty acid synthesis, pertaining to Figure 5. **(A)** Maximum projections of confocal images of calnexin staining in cells treated as indicated (images are from Figures 5B-5C). Scale bar 20 μm. Yellow, cell outlines as determined by overlay and outline drawing over high-brightness ER signal. **(B)** Plot, percent area quantification of ER signal segmentation with phenotypic characterization for comparison. Values per cell (n) and phenotypic categorization are taken from the same dataset as Figures 5B-5C and S6A. **(C)** Plot of incidences of the indicated phenotypes in cells treated with DMSO and 100 µM 1:2:1 palmitic:oleic:linoleic acid or carrier. Data are from the same dataset as Figures 5B-5C and S6A, including DMSO control from 5C. Means + SD shown. P value, Fisher’s exact test.

## STAR Methods

### RESOURCE AVAILABILITY

#### Lead Contact

Requests for further information, resources, and reagents should be directed to Shirin Bahmanyar.

#### Materials Availability

Materials generated in this study are available upon request to the Lead Contact.

#### Data and Code Availability

Raw data generated in this study are available upon request to the Lead Contact.

## EXPERIMENTAL MODEL AND SUBJECT DETAILS

### Mammalian cell lines

U2OS, HEK293, and RPE-1 cells and derived cell lines (see Key Resource Table) were obtained from ATCC or the source specified. U2OS Sec61β-GFP cells were obtained from the Rapoport lab. Cells were grown at 37°C in 5% CO2 in DMEM low glucose (Gibco 11885) (U2OS), DMEM high glucose (Gibco 11965), or DMEM:F12+HEPES (Gibco 113300) supplemented with 2 mM L-glutamine (Sigma 59202C) (RPE-1), all supplemented with 10% heat inactivated FBS (F4135) and 1% antibiotic-antimycotic (Gibco 15240112) or 50 IU/ml penicillin/streptomycin (Gibco 15140). Cells were used for experiments before passage 30 (20 for RPE-1). Cells were tested for mycoplasma upon initial thaw and generation of new cell lines (Southern Biotech 13100-01), and untreated cells were continuously profiled for contamination by assessment of extranuclear DAPI/Hoechst 33258 staining.

### CRISPR/Cas9 genome editing

All guide RNA sequences were designed using the online CRISPR tool http://crispr.mit.edu and reported no off-target matches. CTDNEP1^KO^: ATGAAGTCAGGAGGCGTACC. The guide RNA sequences were synthesized as two oligos with BbsI overhangs and an additional guanidine base 5’ to the protospacer sequence, and the oligos were phosphorylated with calf alkaline intestine phosphatase (New England BioLabs #M0290) and annealed by heating to 95°C and cooling to room temperature. The annealed oligos were cloned into pSPCas9(BB)-2A-Puro (PX459) v2.0 (a gift from Feng Zhang, Addgene plasmid #62988) that had been digested with BbsI-HF (New England BioLabs #R3539). The vector was transfected into U2OS cells using Lipofectamine 2000 and selected with 3 μg/ml puromycin (Invitrogen) for 48 hours. The remaining cells were grown up and gDNA isolated from the bulk population using a QiaAmp DNA Mini kit (Qiagen 51304). Genotyping was performed by sequencing and screening for indels using TIDE deconvolution (https://www.deskgen.com/landing/tide.html; (Brinkman et al., 2014)). Once indels were detected in the bulk population, the cells were plated at <100 cells/ml into 96 well plates and grown in antibiotic-free DMEM with 10% FBS for 2 weeks. Colonies were grown in 24-well plates until more than 10,000 cells could be harvested for gDNA sequencing and TIDE analysis to genotype for frameshift mutations. The CTDNEP1^KO^ clonal cell line used in experiments showed to have +1 insertions in >80% of alleles and 0% WT alleles as determined by TIDE deconvolution of sequencing data.

### Stable Cell Line Generation

To generate U2OS GFP-Sec61β H2B-mCherry, U2OS GFP-Sec61β were transfected with H2B-mCherry-IRES-puro2v2.0 for 48 hours, then plated into 10 cm dishes at <100 cells/ml and selected with 0.5 μg/ml puromycin for 2 weeks. Colonies were trypsinized and picked with 1/8 in sterile cloning discs (Bel-Art F37847-0001) and grown to confluence in a T25 flask before imaging confirmation of marker expression. U2OS CTDNEP1^KO^+CTDNEP1-HA stable cell lines were generated by retroviral transduction, and bulk populations of cells were used for experiments. Retroviruses were generated by transfecting HEK293T cells with pCG-gag-pol, pCG-VSVG and either pMRX-CTDNEP1-HA or pMRX-CTDNEP1-D67ED69T-HA using Lipofectamine 2000. The retroviruses were used to transduce U2OS CTDNEP1^KO^ cells and placed under 7.5 µg/mL blasticidin selection for 2 weeks, then frozen and/or used for experiments. Cells were continuously cultured in 7.5 µg/mL blasticidin.

## METHOD DETAILS

### Transfection and RNAi

Most transfections were performed with Lipofectamine 2000 (Thermo Fisher Scientific 11668) in Opti-MEM (Gibco 31985) using a 1:2 ratio of DNA:lipofectamine with DNA concentrations ranging from 0.05-0.3 μg DNA per cm^2^ of growth surface. Briefly, DNA and lipofectamine were added to 10 μl OptiMEM per cm^2^ of growth surface in separate borosilicate glass tubes (Thermo Fisher Scientific STT-13100-S). After 5 minutes incubation, DNA solution was added to lipofectamine solution. After 15 minutes, DNA:lipofectamine mix was added dropwise to cells plated 16-24 hrs prior to transfection in fresh antibiotic-free media (1 ml/9.6 cm^2^ growth surface). Media was exchanged for antibiotic-free media after 6 hours. Cells were imaged or processed after 24-48 hours. To increase transfection efficiency, plasmids used for live imaging were purified using the Zymopure II Plasmid Midi Prep kit, including a 10 min final elution at 56°C and use of the Zymopure endotoxin removal columns.

Transfections for lipin 1 overexpression were performed using PolyJet in vitro DNA transfection reagent (Signagen SL100688) using a 1:3 ratio of DNA:Polyjet using 0.1 μg DNA per cm^2^ of growth surface. Protocol is identical to previous transfection protocol except for using 5 μl High glucose DMEM per cm^2^ of growth surface for mixes and no incubation before mixing reagents.

For experiments involving transient CTDNEP1 and/or NEP1R1 overexpression, pcDNA3.0 was used as an empty vector negative control. For experiments involving phenotype rescue with transient FLAG-lipin 1β construct overexpression, GFP-KDEL was used as a co-transfection marker, and untransfected cells within the same experiment were used as a negative control for effects of lipin 1β overexpression.

RNAi was performed using Dharmafect 1 (Horizon Discovery T-2001) in Opti-MEM according to the manufacturer’s protocol at the indicated concentrations and durations. RNAi knockdown efficiency was determined with qRT-PCR analysis. U2OS Sec61β/H2B-mCherry were treated with 40 nM CTDNEP1 single siRNA or Ambion Silencer negative control 1 for 48 hours; all others were treated with 20 nM CTDNEP1 siGENOME SMARTpool or control pool siRNA for 72 hours.

### Lipidomics

Early-passage cells were counted by hemocytometer, suspended in PBS at a concentration of 3×10^6^ cells/ml, and flash frozen in liquid nitrogen. Triplicate samples were submitted for each condition, and corresponding triplicate samples were lysed and protein extracted and protein concentration determined by Pierce BCA assay. Sample processing and lipidomics were performed and obtained at Lipotype GmbH. Samples were spiked with lipid class internal standards, and lipids were extracted using chloroform-methanol extraction using a Hamilton Robotics STARlet. Samples were infused using an Advion Triversa Nanomate automated nano-flow electrospray ion source with positive and negative ion mode utilized. Mass spectra were acquired using a Thermo Scientific Q-Exactive hybrid quadruple/Orbitrap mass spectrometer in MS-only or MSMS mode. Lipid species were identified using LipotypeXplorer, and data was processed using Lipotype LIMS and LipotypeZoom. Lipid class pmols/mg protein was determined using protein concentration and sample volume analyzed from each replicate.

### Measurement of Lipin PAP Activity

PAP activity was measured by release of phosphate from [^32^P]PA using Triton X-100 micelles with cell lysates as previously reported (Boroda et al., 2017). Briefly, radiolabeled [^32^P]PA substrate was prepared by phosphorylating 1,2-Dioleoyl-*sn*-glycerol with *E. coli* diacylglycerol kinase and [γ-^32^P]ATP and purified by thin-layer chromatography as described by Han and Carman (DOI: 10.1385/1-59259-816-1:209). To prepare the micelles, Triton X-100 was mixed with buffer A (50 mM Tris-HCl, 10 mM 2-mercaptoethanol, pH 7.4) to a final concentration of 10 mM. Next, 1 μmol of unlabeled 1,2-Dioleoyl-*sn*-glycero-3-phosphate was dissolved in chloroform and mixed with [^32^P]PA (3,000 cpm/nmol) in a glass tube, dried to a thin film under N2 gas, and resuspended with 1 mL of 10 mM Triton X-100. Lysates prepared from U2OS cells containing 10 μg of total protein, radioactive micelles, and buffer A were combined to a final volume of 100 μL, and the reactions were allowed to proceed for 20 min at 30 °C with gentle agitation and were terminated with the addition of 500 μL of acidified methanol (MeOH·0.1N HCl). The final concentrations for all reactions were as follows: 50 mM Tris-HCl, 10 mM 2-mercaptoethanol, and 0.2 mM PA. Free phosphate was extracted with the addition of 1 mL chloroform followed by 1 mL 1M MgCl_2_. The organic extraction was vortexed and 500 μL of the aqueous phase was transferred to a scintillation vial to measure the removal of ^32^P from PA by a scintillation counter. The measurement of PAP activity was determined by following the release of the radiolabeled phosphate from [^32^P]PA. Total PAP activity was measured by including 0.5 mM MgCl_2_. PAP activity for Mg^2+^-independent enzymes was measured by instead including 1 mM EDTA. The activity from assays containing lysate was normalized to activity in assays without enzymes present. Mg^2+^-dependent PAP activity was calculated by subtracting the mean Mg^2+^-independent activity from the total PAP activity.

### ^14^C acetate labeling and lipid extraction

U2OS cells were plated in 24 well plates at a density of 60,000 cells/well and cultured in low-glucose DMEM (Gibco, 11885-084) with 10% FBS (Gemini Bio-products, 900-108) and 1% antibiotic-antimycotic (Gibco, 15240-062). Cells were labeled with ^14^C acetate and lipids extracted to determine acetate incorporation into neutral lipids (Liebergall et al., 2020). 16-18 hours after plating, cells were labeled with 500 μl fresh media containing 1 μCi/mL (19.23 μM) 1,2-^14^C-acetate (Perkin-Elmer, NEC553) for 5 hours. Cells were then washed with PBS 2x on ice, lysed with 250 μL of 0.1% Triton X-100, 0.5 mM DTT, and a protease inhibitor cocktail (10 μg/mL leupeptin, 10 μg/mL pepstatin A, and 1 mM phenylmethylsulfonyl fluoride) in PBS, pH 7.2, and homogenized by pushing through a 22G needle 6x. 200 μl of lysate was added to 500 μl 0.1 N HCl·methanol, then 250 μl chloroform was added. The extract was vortexed and incubated for 1-2 min at room temp. Another 250 μl of chloroform was added, followed by 250 μl of 0.2M NaCl. The extract was vortexed and centrifuged at 1000xg for 1 min. The aqueous phase was then aspirated, and 250 μl of the organic phase was used for quantification of ^14^C by a Beckman-Coulter scintillation counter. To calibrate counts per minute per nmol ^14^C-acetate, 250 μl media containing 1 μCi/ml ^14^C acetate was counted. Remaining cell lysates were used for BCA protein quantification for normalizing to protein concentration. To account for background ^14^C quantification, a control set of cells was treated with 10 μM 5-(tetradecyloxy)-2-furoic acid (TOFA) in DMSO 30 min before and during ^14^C labeling, and these values were subtracted from final values and shown separately. Results were expressed as nmol^14^C acetate incorporated into lipids per mg of protein.

### Fatty acid supplementation

Cells were plated at a density of 200,000 cells/ml in 6 well plates. Stocks of oleic acid, linoleic acid, and palmitic acid were made in methanol and pipetted into a 50 ml conical vial, then dried with an ambient air stream. Pre-warmed DMEM containing 0.5% fatty acid-free BSA (Sigma Cat#A8806) was added to a final concentration of 25 μM palmitic acid, 50 μM oleic acid, and 25 μM linoleic acid (100 μM total fatty acid concentration; 1:2:1 ratio of palmitic:oleic:linoleic acid). The solution was incubated at 37°C for 30 min, then held to the bottom of a sonicating bath for 30 s, then incubated at 37°C for 10 min until solution was clear. FBS was added to a final concentration of 10% v/v. Cells were treated with DMEM with 10% FBS and 0.5% BSA alone or DMEM with 10% FBS 0.5% BSA, and 100 μM fatty acids with DMSO or 10 μM TOFA in DMSO for 24 hrs prior to immunofluorescence processing.

### Mitotic and micronuclei enrichment

For mitotic shakeoff to enrich for M phase cells, cells were grown to at least 50% confluence in 75 cm^2^ flasks. Cells were washed with PBS or antibiotic-free media to clear debris, then flasks were whacked repeatedly on all sides and tapped on the bottom surface with a reflex hammer (DR Instruments S72118) until at least 50% of mitotic cells were dislodged. The cell media was collected and centrifuged at 300xg for 5 min, then cells were additionally washed or plated.

For scoring of intracellular ER membranes in prometaphase-metaphase cells, whole 75 cm^2^ flasks of cells were transfected with GFP-KDEL and imaged 48 hours later after mitotic shakeoff and plating into 1 well per flask of an ibidi 8-well imaging chamber. Cells expressing GFP-KDEL in prometaphase up until metaphase (determined by DIC chromatin appearance) were imaged with 0.5 μm stacks for 20 μm total z height.

For enrichment of cells at the G2-M phase transition, cells were treated with 9 μM RO-3306 for 17-20 hours (Vassilev, 2006). On the microscope stage for live imaging, media was exchanged for Fluorobrite DMEM with 10% FBS and 1 mM L-glutamine 7 times, then imaging was initiated within 5-10 min of the first wash.

For micronuclei enrichment using nocodazole washout, cells at 50-80% confluence in 75 cm^2^ flasks were washed with 37°C PBS to clear debris and then treated with 100 ng/ml nocodazole (Sigma M1404) in antibiotic-free media for 6 hours (Liu et al., 2018). Cells were subject to mitotic shakeoff without washing, then washed 3x with 37°C PBS. After the final wash, cells were plated onto acid-washed coverslips (coated with 1 μg/ml poly-D-lysine (Sigma P7886) for short-term washout) and incubated for 45 min-60 min (short-term) or 18-20 hours (long-term) before immunofluorescence processing.

For RO-3306/MPS1i micronuclei enrichment, cells were treated with 9 μM RO-3306 (Calbiochem 217699) for 19 hours, then with 1 μM NMS-P715 (MPS1i) (Calbiochem 475949) for 18 hours before processing for immunofluorescence (Liu et al., 2018).

### Quantitative real-time PCR

RNA was harvested using the RNeasy Mini kit (Qiagen 74104) using the manufacturer’s protocol, using Qiashredders (Qiagen 79654) for tissue homogenization and with additional RNase-free DNase (Qiagen 79254) treatment after the first RW1 wash and subsequently adding another RW1 wash. RNA was eluted with RNAse-free water and diluted to 50 ng/μl. RNA was subject to reverse transcription using the iScript Reverse Transcription Supermix (Bio-Rad 1708840) with 400 ng RNA per reaction. The subsequent cDNA was diluted 1:5 for RT-qPCR. cDNA was analyzed for RT-qPCR using the iTaq universal SYBR Green Supermix (Bio-Rad 1725120). Cycle threshold values were analyzed using the ΔΔCt method.

### Immunofluorescence

Cells were washed 2x with warm PBS and fixed in 4% paraformaldehyde (+0.1% glutaraldehyde for ER structure analyses) in PBS for 15 min, permeabilized in 0.5% Triton X-100 for 5 min, then washed 3 times with PBS and blocked in 3% BSA in PBS for 30 min. Samples were transferred to a humidity chamber and incubated with primary antibodies in 3% BSA in PBS for 1 hour at room temperature with rocking. Samples were washed with PBS 3 times for 5 min, then incubated with secondary antibodies in 3% BSA in PBS for 1 hour at room temperature in the dark with rocking. Samples were then washed with PBS 3 times for 5 min in the dark. For experiments visualizing nuclear structure and/or micronuclei, cells were additionally stained with 1 μg/ml Hoechst 33258 (Thermo Fisher Scientific H3569) in PBS for 1 min followed by one PBS wash. Coverslips were mounted with ProLong Gold Antifade reagent + DAPI (Thermo Fisher P36935) and sealed with clear nail polish. For samples treated with goat primary antibodies, 5% normal donkey serum (Sigma D9663) was used in place of 3% BSA.

When indicated, cells were fixed and stained to visualize kinetochore microtubules (Thompson and Compton, 2011) by extracting in 100 mM PIPES, 1 mM MgCl_2_, 1 mM CaCl_2_, 0.5% Triton X-100, pH 6.8 for 4 min, then fixing in 1% glutaraldehyde in PBS for 10 min and quenched 2 times with 0.1% NaBH_4_ in TBS for 10 min each. Cells were washed twice with 10 mM Tris, 150 mM NaCl, 10% BSA and then stained with tubulin antibody for 1.5 hours, washed with PBS, then stained with secondary antibody for 1 hour, washed with PBS, then mounted with ProLong Gold + DAPI. Antibody concentrations used were: Rabbit anti-calreticulin 1:100; Rabbit anti-emerin 1:200; Mouse anti-α tubulin DM1A 1:1000; Mouse anti-FLAG 1:1000; Rabbit anti-HA 1:800; all secondaries, 1:200-1:250.

### Immunoblot

Lysis buffers used were: RIPA buffer (1% NP-40, 0.5% sodium deoxycholate, 0.1% SDS, 150 mM NaCl, and 1 tablet/50 ml cOmplete protease inhibitor cocktail in 25 mM Tris pH 7.4); or 0.1% Triton X-100, 50 mM NaF, 1mM EDTA, 1 mM EGTA, 10 mM Na_2_HPO_4_, 50 mM β-glycerophosphate, 1 tablet/50 ml cOmplete protease inhibitor cocktail, pH 7.4 (for lipin 1 analysis). Cell lysates were removed from growth surfaces by scraping with a rubber policeman after incubation in lysis buffer or by adding lysis buffer to cell pellets collected by trypsinization and centrifugation at 300xg for 5 min followed by 1-2 PBS washes. Lysates were homogenized by pushing through a 23G needle 30 times and then centrifuged at >20,000xg for 10 min at 4°C, then protein concentration was determined using the Pierce BCA Protein assay kit (Thermo Scientific 23225). 10-30 μg of whole cell lysates/lane were run on 8-15% polyacrylamide gels dependent on target size, and protein was wet transferred to 0.22 μm nitrocellulose (<100 kDa) or PVDF (>100 kDa) membranes. Ponceau S staining was used to visualize transfer efficiency, then washed with TBS or DI water; then, membranes were blocked in 5% nonfat dry milk or BSA in TBS for 1 hour. Membranes were then incubated with primary antibodies in 5% milk or BSA for 1-2 hours at room temperature or overnight at 4°C with rocking. Membranes were washed 3 times for 5 min in TBS-T, then incubated with anti-HRP secondary antibodies in 5% milk or BSA in TBS-T for 1 hour at room temperature with rocking. Membranes were washed 3 times for 5 min in TBS-

T. Clarity or Clarity Max ECL reagent (Bio-Rad 1705060S, 1705062S) was used to visualize chemiluminescence, and images were taken with a Bio-Rad ChemiDoc or ChemiDoc XRS+ system. Exposure times of images used for analysis or presentation were maximum exposure before saturation of pixels around or within target bands. Antibody concentrations used were: Mouse anti-α tubulin DM1A 1:5000; Mouse anti-FLAG 1:4000; Rabbit anti-HA 1:1000; Rabbit anti-lipin 1 1:1000; Rabbit anti-Phospho-p70 S6 Kinase (Thr389) (108D2) 1:1000; Rabbit anti-p70 S6 kinase 1:1000; Rabbit anti-Phospho 4E-BP1 (Ser65) 1:1000; Rabbit anti-4E-BP1 1:1000; all secondaries 1:10000

### Live cell imaging

For live imaging, cells were plated in Willco Wells 35 mm dishes (Willco Wells HBST-3522), ibidi 2 well imaging chambers (ibidi 80287) with DIC lid (ibidi 80055); ibidi 8 well imaging chambers (ibidi 80827). Samples were imaged in a CO2-, temperature-, and humidity-controlled Tokai Hit Stage Top Incubator. Objectives were also heated to 37°C. For CO2-controlled imaging, the imaging media used was Fluorobrite DMEM (Gibco A1896701) supplemented with 10% FBS. U2OS GFP-Sec61β/H2B-mCherry mitotic cells were imaged using a custom aluminum stage insert heated to 37°C with heating tape and temperature monitored using a Physitemp thermistor (BAT7001H) and probe (IT-18), with objective heating and using 140 mM NaCl, 2.5 mM KCl, 1.8 mM CaCl_2_, 1.0 mM MgCl_2_, 20 mM HEPES, 15 mM glucose, pH 7.4 as the live cell imaging solution. When indicated, cells were treated with 1 μM SiR-DNA (Cytoskeleton, Inc. CY-SC007) for 1 hour prior to imaging and kept in SiR-DNA-containing live imaging media during imaging. When indicated, cells were treated with 1 μM ER Tracker Green (Invitrogen E34251) for 30 min and washed out prior to imaging, and cells were imaged for a maximum of 2 hours after treatment.

### Cytoplasmic viscosity measurements

To prepare for cytoplasmic viscosity measurements, 2.75+/-0.219 μm diameter streptavidin-functionalized superparamagnetic beads (Bangslabs PMS3N) were incubated with 1 μM 647 fluorophore (Atto647N Biotin, Sigma 93606) for 2 hours at 4 C, shaking at 400 RPM. Cells were plated with beads during passaging and left to grow for 2-3 days until 75-80% confluent. Cells were passaged in this way for 3-4 cycles until visual inspection of cells shows approximately 40% of mitotic cells contain a single bead. Cells were grown in 35 mm Desag 263 glass-bottomed culture dish (0.17 mm thick glass black DeltaT, 04200417B, Bioptechs) to 75-80% confluency Prior to experiments, cell medium was switched to an imaging medium (FluoroBrite DMEM, Gibco) supplemented with 10 mM HEPES and left to equilibrate for 10-20 minutes. Then a layer of 1 mL white mineral oil (VWR) was added on top of the imaging medium, and the dishes were mounted in a temperature control system and kept at 37 °C (DeltaT Culture Dish System, Bioptechs). Only metaphase cells displaying proper chromosome alignment, no significant blebbing or morphological issues, and a high expression level of the expected tags, and having a single bead were selected for experiments.

The cylindrical magnetic tweezer core was made of ¼” wide, 6” long HyMu80 alloy (EFI Alloy 79, Ed Fagan Inc.) and sharpened at one end to a cone with a tip width of 5 μm. The solenoid frame was a steel cylinder 1” wide, 3” long, and has a hole for the core to fit through. The frame was held onto the core using 2 set screws. The solenoid was made using sheathed, 24-gauge copper wire (7588K77, McMaster-Carr) and wound 400 times. The solenoid and core were mounted on a micromanipulator (NMN-21, Narishige), which was mounted on a custom-built base using ThorLabs components. The solenoid was connected to a programmable power supply (PSP-603, GW Instek).

To analyze cytoplasmic viscosity measurements, we built a custom MATLAB pipeline. Briefly, we used featuretrack, an open-source MATLAB package by Maria Kilfoil, modified with a 3D Gaussian fit to localize the bead each frame and calculate displacement between successive frames. Once we found the full trajectory of the bead, we fit the velocity, bootstrapping using 50% of the data, repeated 10 times. We had previously calibrated the magnetic tweezer system using the same beads suspended in glycerol as described (Kollmannsberger and Fabry, 2007) to produce a 2D map of the force exerted on the bead based on the location relative to the tweezer tip. To calculate cytoplasmic viscosity, we rearranged Stokes’ Law, which defines the drag force on a sphere moving through a viscous fluid:

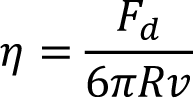

where F_d_ is the drag force, *R* is the bead radius, *v* is the bead velocity, and η is the effective cytoplasmic viscosity for particles similar in size to the magnetic bead. Final reported data points represent a calculation of cytoplasmic viscosity per pull across multiple cells. Errors were calculated taking into account tweezer calibration, bead localization, and the velocity fit.

### Microscopy

Samples were imaged on an inverted Nikon Ti microscope equipped with a Yokogawa CSU-X1 confocal scanner unit with solid state 100-mW 488-nm and 50-mW 561-nm lasers, using a 60×1.4 NA plan Apo oil immersion objective lens, and a Hamamatsu ORCA R-2 Digital CCD Camera.

Samples with SiR-DNA/GFP-KDEL or FLAG-lipin/calnexin staining or telophase nocodazole washout cells were imaged on an inverted Nikon Ti Eclipse microscope equipped with a Yokogawa CSU-W1 confocal scanner unit with solid state 100 mW 405, 488, 514, 594, 561, 594, and 640 nm lasers, using a 60x 1.4 NA plan Apo oil immersion objective lens and/or 20x plan Fluor 0.75 NA multi-immersion objective lens, and a prime BSI sCMOS camera.

Samples for magnetic tweezer experiments and chromosome velocity measurements were imaged on an inverted Nikon Ti Eclipse equipped with a manual rotation stage. Multi-dimensional time-series images were acquired with a Yokogawa CSU-X1 spinning disk unit with a 1.2X camera mount magnifier, Coherent Obis lasers (488, 5660, 640 nm), a motor-driven filter wheel (filters: 514/60, 593/40, 647 LP; Ludl), an objective z-piezo stage (Physik Instrumente), a 60x 1.4 NA plan Apo oil immersion objective lens, and a sCMOS camera (Flash LT+, Hamamatsu).

### Image analysis

Image analysis was performed using FIJI/ImageJ unless otherwise noted. For scoring of ER phenotypes, cells expressing moderate levels of GFP-KDEL with no overexpression artifacts (dense fluorescent clumps in ER or nuclei) were included for analysis. For scoring of interphase ER expansion, cells with a network of peripheral ER tubules visualized with GFP-KDEL or calnexin staining were considered “normal”, while cells with ER sheets and tubules extending into the periphery with a lack of any tubular network were considered to have “expanded ER”. Additionally, cells with the appearance of thin ER tubules, large gaps between tubules, and a smaller cluster of perinuclear ER were considered to have “reduced ER” with TOFA treatment. These phenotypes were additionally quantified with percent abundance of cytoplasmic KDEL/calnexin signal: for cells with the entire ER captured within 0.3-0.5 μm interval z stacks, 8 bit maximum intensity projections were made of the whole field of view. To ensure the different ER morphologies were all accounted for after thresholding, the 8-bit max projections were subject to unsharp masking with a radius of 2 and mask of 0.6. The max intensity projection was thresholded using the Huang threshold of object fuzziness (Huang and Wang, 1995). The cell border and nuclear border for each cell were manually traced using ER fluorescent signal, and the percent of KDEL-positive pixels per nucleus-free cell area was measured. RPE-1 cell phenotypes were scored blindly.

For quantification of micronuclei, images taken at 60x were scored for presence of micronuclei (DNA fragments encased in an emerin or calnexin-positive rim apart from main nucleus <∼20% in size of the main nucleus). Severely lobulated/partitioned “hypermicronucleated” nuclei (DNA fragments/lobes apart from the main nucleus >∼20% in size of the main nucleus) and micronuclei were scored through oculars or in 60x images of cells with nuclear envelope staining. Nuclei with both lobes/partitions and micronuclei were considered hypermicronucleated. For quantification of peripheral chromosome/tubulin masses in cells subjected to short-term nocodazole washout, 60x images of cells processed for immunofluorescence without non-kinetochore microtubule depolymerization were scored for the presence of chromosome masses with microtubules extending to them that were to away to the cell periphery compared to the primary nuclei chromosome masses.

Nuclear solidity was quantified as described (Fonseca et al., 2019). Briefly, DAPI/Hoechst images were thresholded with the ImageJ default setting, then the magic wand tool was used to select segmented nuclei. Nuclei that were unable to be segmented due to poor signal:noise, adjacent nuclei touching, or presence of a micronucleus touching the main nucleus were not included in the analysis. Segmented and selected nuclei were measured using the ImageJ shape descriptors measurement metric. Data were expressed as % of nuclei with a solidity value less than the control U2OS average minus 1 standard deviation.

To quantify the percent of mitotic cell diameter that is occupied by ER membranes in cells expressing GFP-Sec61β/H2B-mCherry or GFP-KDEL/SiR-DNA, 60x image stacks of cells at anaphase onset (determined by first frame of visible chromatid separation) were obtained. Image background was subtracted using the average value of 3 boxes from surrounding the cell (but not within adjacent cells). A 10-pixel thick line was drawn encompassing the cell diameter along the metaphase plate (in the center of the dividing chromatin masses, along the division plane), and a profile plot was generated. The local maxima of the Sec61β/KDEL peaks for each side of the cell was determined, and the width of the half maxima for each of the 2 Sec61β/KDEL peaks was quantified and added together. This value was divided by the diameter of the cell (determined by the bounds of the Sec61β/KDEL half maxima) to determine the % of the cell diameter occupied by ER signal. For representation, plot profiles shown are normalized to minimum and maximum of ER and DNA signal.

For quantification of intracellular ER membranes in prometaphase-metaphase cells (determined by DIC chromosome appearance), 90x images of cells expressing GFP-KDEL and subject to mitotic shakeoff were blindly categorized for presence of a) no intracellular ER membranes (“cleared”), (b) few ER tubules within the cell interior (“partially cleared”); or c) large (>2 μm length) sheets and/or several tubules within the cell interior (“not cleared”).

To analyze average chromosome velocity in prometaphase cells, we used PIVlab, an open-source MATLAB toolbox for particle image velocimetry, to generate velocity fields of chromosome movement within the chromosome mass. We then used MATLAB to filter and analyze velocity information. Final reported chromosome movements correspond to the average velocity magnitude of the chromosome mass deformation per frame averaged over time.

To measure nuclear:cytoplasmic ratio of lipin expression, 3 10×10 pixel regions in the nucleus and cytoplasm from the same focal plane were taken and average intensities measured. These 3 average intensities were then averaged and the value for cytoplasmic signal divided by the value for nuclear signal. Background was subtracted using a region outside of the cell.

### Statistical analysis

GraphPad Prism 8 was used for all statistical analysis. Continuous data was tested for normality using a Shapiro-Wilk test. For experimental setups in which > 10 samples (n) per experimental replicate (N) were able to be collected consistently, continuous data was measured with paired t tests of experimental replicate means. Superplot format was used for representing percent of ER-positive pixels in cytoplasm area (Lord et al., 2020). Experimental replicates of discrete data were plotted with shapes indicating separate replicates to display reproducibility, and incidences between groups (replicates pooled) were tested for significance using Fisher’s exact test (2 categories) or Chi square test (>2 categories). Statistical tests used, sample sizes, definitions of n and N, and p values (p<0.05 as significance cutoff) are reported in figures and/or figure legends. For quantification of all data where >10 samples could be gathered within an experimental repeat, sample size calculations using the online tool (https://clincalc.com/stats/samplesize.aspx) determined the adequate sample size for number of cells to analyze for sufficient (80%) power.

## Movie Legends

**Movie S1. ER membrane clearance and dynamics during anaphase and mitotic exit, pertaining to** **Figure 2**. Confocal time lapse images of GFP-Sec61β (gray)/H2B-mCherry (magenta) in U2OS cells treated with control (left) or CTDNEP1 (right) siRNA. Time stamp (min:sec). Scale bar 10 µm.

**Movie S2. Unaligned chromosomes embedded in ER membranes during prometaphase, pertaining to** **Figure 2**. Confocal time lapse images of GFP-KDEL (gray) and H2B-mCherry (magenta) in transiently-expressing control (left) and CTDNEP1^KO^ (right) U2OS cells taken after Cdk1i washout. Arrows point to unaligned chromosomes during prometaphase. Scale bar 10 µm.

**Movie S3, Mitotic cytoplasm viscosity measurements taken from displacement of beads upon application of constant force with magnetic tweezers, pertaining to** **Figure 2**. Confocal time lapse images of ER Tracker Green staining (gray) in U2OS cells cultured with Alexa Fluor 647-conjugated 2.75 µm magnetic beads (magenta). Beads are pulled through the cytoplasm using magnetic tweezers. Time stamp (min:sec). Scale bar 10 µm.

**Movie S4, Prometaphase chromosome movements, pertaining to** **Figure 2** Confocal time lapse images SiR-DNA staining in control (left) and CTDNEP1^KO^ (right) U2OS cells. Time stamp (min:sec). Scale bar 10 µm.

**Table S1, Supplement to the Key Resources Table**

